# Disruption of *Zea mays isochorismate synthase1* decreases PHENYLALANINE AMMONIA LYASE activity and suppresses hypersensitive response-induced metabolism

**DOI:** 10.1101/2022.11.04.515247

**Authors:** Ryan L. Benke, Rachel M. McCoy, Iskander M. Ibrahim, Jeffery P. Simpson, Fabiola Muro-Villanueva, Ross Zhan, Clint Chapple, Joshua R. Widhalm, Sujith Puthiyaveetil, Gurmukh S. Johal, Brian P. Dilkes

## Abstract

ISOCHORISMATE SYNTHASE (ICS) catalyzes the isomerization of chorismate to isochorismate, an essential precursor in the biosynthesis of the Photosystem I electron carrier phylloquinone and of one of two pathways for the biosynthesis of the defense response hormone salicylic acid (SA). We characterized a *Zea mays ics1* mutant for impacts on metabolism, photosynthesis, and immune signaling. Phylloquinone was reduced in the mutant resulting in low electron transfer rates and high electron backflow rates. SA accumulation induced by autoactive alleles of the nucleotide-binding leucine-rich repeat (NLR) gene *Resistance to Puccinia sorgi1* (*Rp1)* required *ics1*. Induced accumulation of SA was not required for lesion formation by the autoactive *Rp1-D21#4* allele. Metabolomic analyses and SA supplementation of *Rp1-D21#4* mutants, *ics1-1* mutants and *Rp1-D21#4; ics1-1* double mutants demonstrated that most hypersensitive response-induced metabolism required *ics1* but this was independent of SA accumulation. Both the PAL and ICS pathways contributed to SA biosynthesis in maize as labeled phenylalanine was incorporated into SA glucoside. Maize *ics1-1* mutants had low PHENYLALANINE AMMONIA LYASE activity, accumulated phenylalanine, and decreased abundance of phenylalanine derived metabolites. This demonstrates that the ICS and PAL pathways interact by a yet unknown mechanism complicating the interpretation of SA biosynthesis in plants from genetics alone.

## Introduction

ISOCHORISMATE SYNTHASE (ICS) catalyzes the reversible conversion of chorismate to isochorismate (Figure 1), an essential precursor in the biosynthesis of the Photosystem I (PSI) electron carrier phylloquinone (Gross et al., 2006; Garcion et al., 2008; Qin et al., 2019). Plants deficient in phylloquinone are pale, stunted in growth, and exhibit germination defects and seedling lethality (Garcion et al., 2008; Qin et al., 2019; Orcheski et al., 2015; Emonds-Alt et al., 2017). In plants, ICS activity is encoded by the multifunctional gene *PHYLLO* and/or by a stand-alone *ICS* gene. All Viridiplantae contain a *PHYLLO* gene derived from a fusion of genes homologous to bacterial *MenF* (ICS), *MenD* (E.C. 2.2.1.9), *MenC* (E.C. 4.2.1.113), and *MenH* (E.C. 4.2.99.20). In *Physcomitrella patens* and green algae, the tetra-fused PHYLLO is responsible for the first four steps in phylloquinone biosynthesis, including isochorismate synthesis. Angiosperms, however, encode two sub-functionalized duplicates of the multifunctional *PHYLLO* gene. One duplicate encodes the activities of MenD, MenH, and MenC, but a deletion within the MenF derived sequences removed the chorismate binding domain (Gross et al., 2006). The other duplicate, annotated as ICS, retained the MenF derived sequences but has lost the MenD, MenC, and MenH derived sequences and is necessary and sufficient for the biosynthesis of isochorismate (Garcion et al., 2008; Qin et al., 2019).

**Figure 1.**
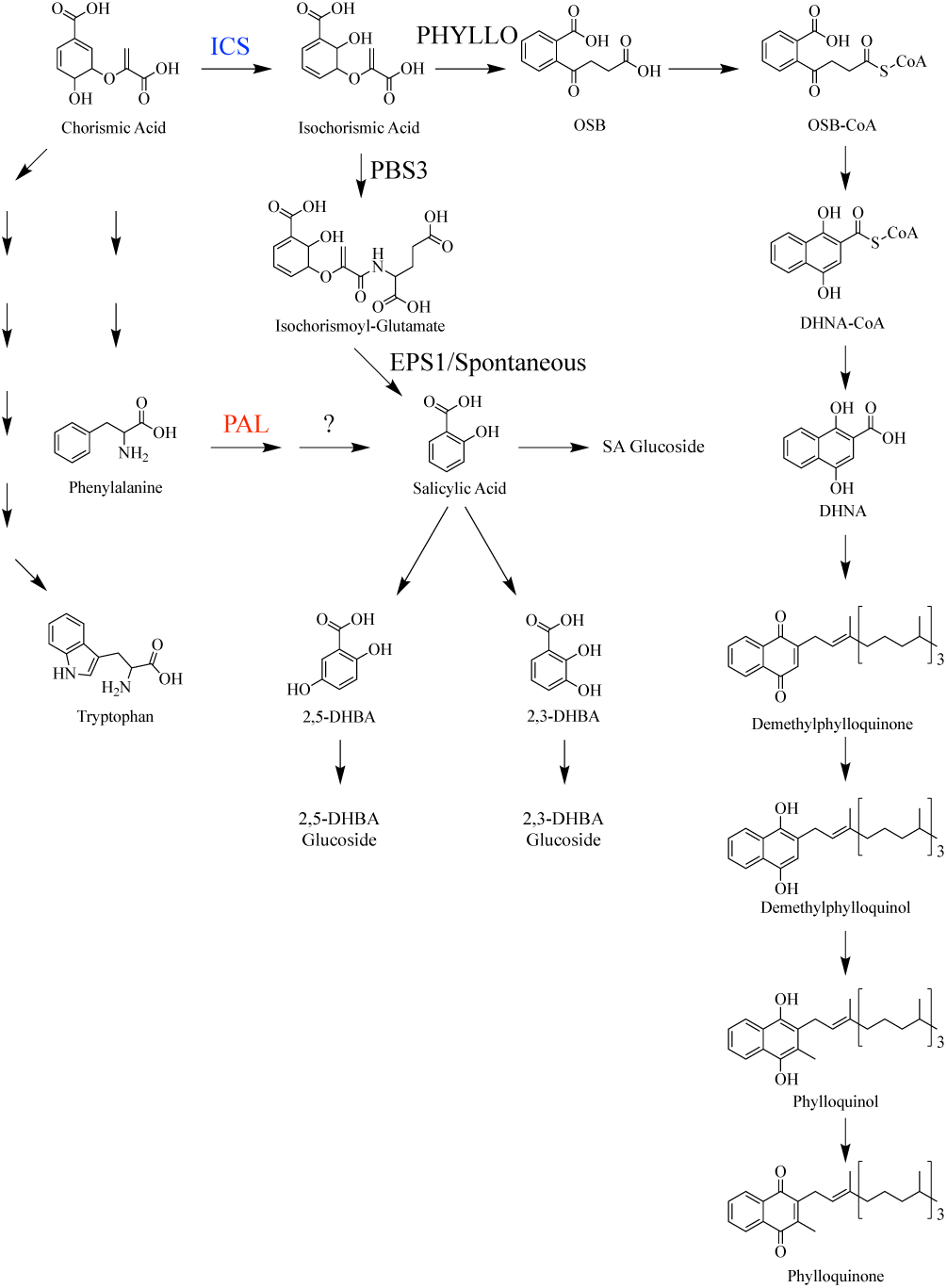
Biosynthetic pathways of phylloquinone and salicylic acid in plants.

Isochorismate is also an intermediate in one of the two biosynthetic pathways for salicylic acid (SA) in plants. In the ICS-dependent SA pathway, isochorismate is converted to isochorismoyl-glutamate by the enzyme encoded by *avrPphB SUSCEPTIBLE3* (*PBS3),* and isochorismoyl-glutamate is spontaneously and/or enzymatically cleaved to form SA (Figure 1; Rekhter et al., 2019; Torrens-Spence et al., 2019). The other route to synthesize SA in plants is from L-phenylalanine (Figure 1). Although the genes encoding all steps of the pathway from phenylalanine to SA have not been identified, both genetic evidence (Huang et al., 2010; Yuan et al., 2019) and isotope-labeling analyses (Pallas et al., 1996; Coquoz et al., 1998) indicate the requirement of PHENYLALANINE AMMONIA LYASE (PAL), which deaminates phenylalanine to *trans*-cinnamic acid. In multiple plant species, benzoic acid also serves as a precursor for SA in the PAL-dependent SA pathway (H. D. KLÄMBT, 1962; Yalpani et al., 1993).

Genetic evidence suggests that SA production from the ICS-dependent and PAL-dependent pathways is unequal. In Arabidopsis, knockout of *ics1* resulted in a greater than 90% decrease in pathogen-induced SA production (Wildermuth et al., 2001; Huang et al., 2010). As such, the PAL-dependent pathway has been treated as a minor contributor to SA biosynthesis in Arabidopsis. However, genetic disruption or virus-induced silencing of *PAL* genes in Arabidopsis, soybean, and maize resulted in a greater than 50% reduction in pathogen-induced SA accumulation (Huang et al., 2010; Shine et al., 2016; Yuan et al., 2019). Thus, these two pathways appear to be interdependent, as the combined decrease in SA accumulation when targeting either the PAL-dependent or ICS-dependent SA biosynthetic pathways in Arabidopsis (Huang et al., 2010) and soybean (Shine et al., 2016) exceeds one hundred percent.

Studies on the function of SA in plants have focused on its role in immune signaling and defense response. SA can act as a signaling molecule perceived by NONEXPRESSOR OF PATHOGENESIS-RELATED GENES1 (NPR1) or by NPR3/4. The binding of SA to NPR1 or NP3/4 results in transcriptional activation of pathogenesis-related genes by two distinct mechanisms (Ding et al., 2018). SA signaling plays functional roles in both effector- and pattern-triggered immunity, and SA is required for both local and systemic responses to numerous pathogens (Vlot et al., 2009). Much of our understanding of SA function in plants comes from work in tobacco and Arabidopsis, where the genetic resources to manipulate SA accumulation are well developed (Gaffney et al., 1993; Lawton et al., 1995; Wildermuth et al., 2001; Garcion et al., 2008). These resources are lacking for most plants, and the functions of SA in plant immunity responses have not been adequately tested in plants other than tobacco and Arabidopsis.

Accumulation of SA is a hallmark of the plant hypersensitive response (HR), the rapid localized cell death at and around the site of pathogen infection (Balint-Kurti, 2019). Studies of HR and HR-induced molecular signaling have been facilitated by lesion mimic mutants encoded by autoactive alleles of resistance (R) genes and display constitutive HR-signaling and lesion formation in the absence of pathogen infection (Bruggeman et al., 2015). These mutants are advantageous for molecular studies of HR, as they eliminate the need to control for pathogen infection, and all measured molecular signals can be attributed to the biology of the plant. In maize, the most well-studied autoactive R-gene mutant is *Rp1-D21*. This mutant originated from an intergenic recombination at the *Rp1* locus, which encodes a set of tandemly repeated nucleotide-binding site leucine-rich repeat (NLR) proteins that mediate resistance to the common rust fungus, *Puccinia sorghi* (Hu et al., 1996)*. Rp1-D21* mutants accumulate high levels of SA (Ge et al., 2021), but a requirement of SA for lesion formation and HR-responsive molecular signaling in *Rp1-D21* mutants has yet to be tested.

In this work, we characterized a loss-of-function allele of the *ICS1* gene of maize and investigated the biosynthesis and function of SA. Using metabolite analysis, labeled isotope feedings, and enzyme-activity assays, we provide evidence for the existence of both the ICS-dependent and PAL-dependent SA biosynthetic pathways in maize. We demonstrate that the two pathways are not independent and that loss of ICS-activity in maize suppresses PAL-activity. This is coincident with increased accumulation of phenylalanine and decreased accumulation of phenylalanine-derived metabolites. Double mutants of *ics1-1* and *Rp1-D21#4* do not accumulate SA but do initiate lesions, demonstrating that SA hyperaccumulation is not required for this phenotype. Metabolite profiling shows that SA accumulation is also dispensable for the differential accumulation of most HR-responsive metabolic perturbations in maize.

## Results

### Characterization of Zea mays isochorismate synthase

The *Zea mays* B73 version 4 reference genome contains a single annotated *ics* gene, which was previously named *isochorismate synthase like1* (*ics1;* Zm00001d020220). The predicted gene product of *ics1* contains a putative 21-amino acid N-terminal chloroplast signaling peptide (Emanuelsson et al., 1999), a chorismate binding domain motif (Pfam accession no. PF00425), and four amino acid residues conserved across all ICS enzymes and required for substrate binding and catalysis (Figure 2; Kolappan et al., 2007; Garcion et al., 2008). To determine if there are any other *ICS*-like genes in maize, we performed tBLASTn searches of the *Zea mays* B73 version 4 reference genome using the exon sequences of maize *ics1* and Arabidopsis *ICS1* and *ICS2* as query. The maize *ics1* gene was the only ICS homolog identified among these BLAST hits, indicating that the maize genome encodes a single isochorismate synthase gene.

**Figure 2.**
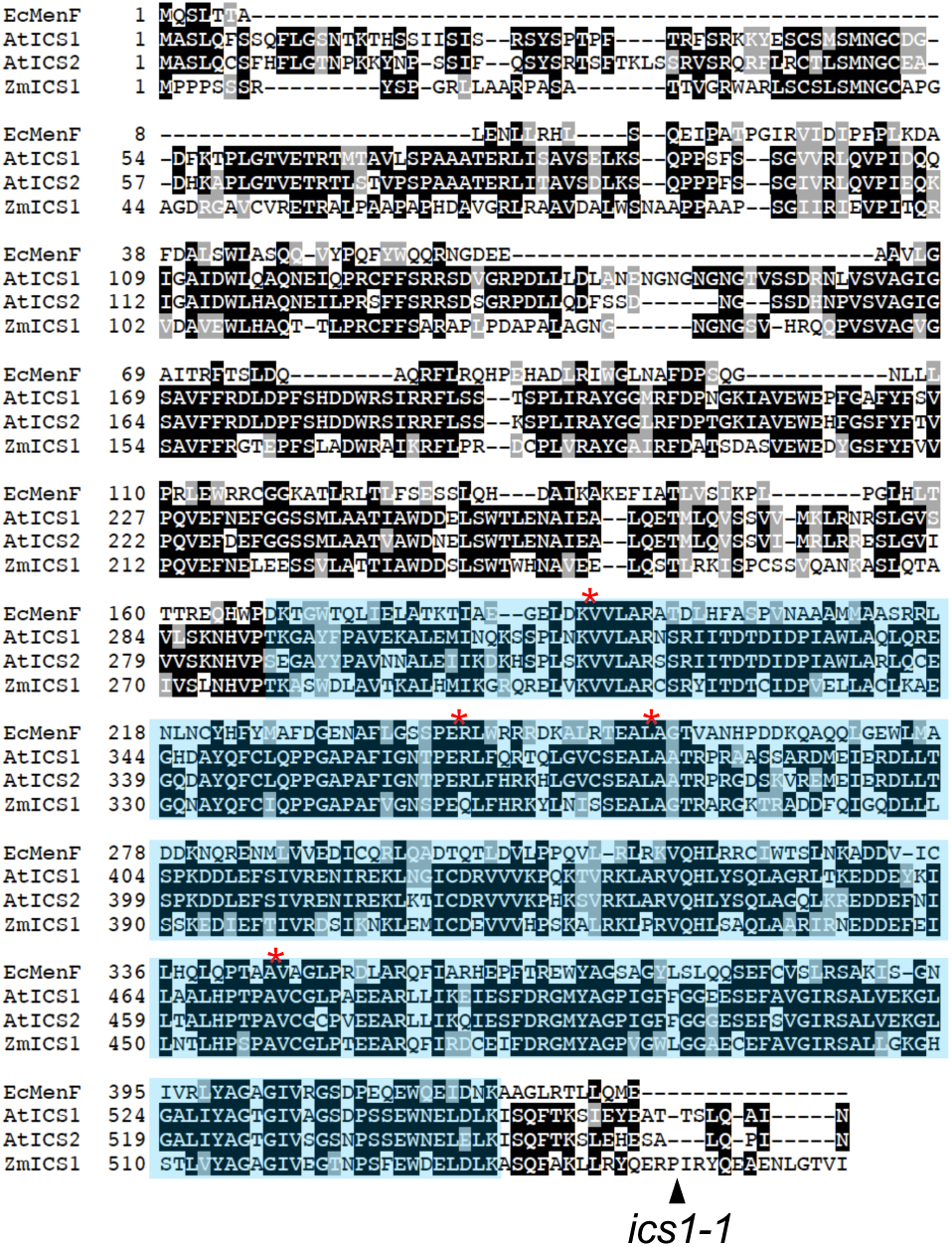
*Zea mays* encodes a single *isochorismate synthase* gene. Sequence alignment of the amino acid sequence of *Escherichia coli* MenF, *Arabidopsis thaliana* ICS1 and ICS2, and *Zea mays* ICS1. Conserved lysine (*E. coli* K190), glutamic acid *(E. coli* E240), leucine *(E. coli* L255), and alanine (*E. coli* A344) residues required for ICS activity are denoted by red stars. The highly conserved chorismate binding domain (Pfam accession no. PF00425) is shaded in blue. The putative 21 amino acids that make up the maize ICS1 chloroplast targeting peptide are highlighted in green. Mutator insertion position for *ics1-1*, relative to the *Z. mays* ICS1 amino acid sequence, is labeled by a black arrowhead.

Previously characterized plant *ICS* genes have 13, 14, or 15 exons with 15 exons representing the most likely ancestral *ICS* gene structure (Yuan et al., 2009). Maize *ics1* contains 14 exons and has lost intron two compared to the 15-exon structure (Figure 3). To determine when plant *ICS* genes underwent exon-intron boundary modifications and any gene duplications, a phylogenetic analysis of angiosperm *ICS* genes was overlayed with the gene structure changes and gene duplication events (Figure 3). Two *ICS* paralogs are present in *Brassicaceae* consistent with a duplication at the base of this lineage. Lineage-specific duplications, consistent with recent polyploidy events, are present in *Glycine max*, *Gossypium raimondii*, and *Arachis hypogaea*. The differences in gene structures of the *ICS* genes included in this phylogeny could be explained by four intron-loss events, relative to the 15-exon structure. The loss of intron five, a feature that distinguishes the Arabidopsis *ICS1* and *ICS2* paralogs, occurred within *Brassicaceae* after the divergence of *Eutrema salsugineum* from the other species analyzed. The *ICS1* gene from Arabidopsis has further undergone a lineage-specific loss of intron three. Intron two was lost twice, once at the base of the *Fabaceae* lineage and once within the panicoid grasses. The 14-exon maize *ics1* gene structure is consistent with the gene structure present in other panicoid grasses.

**Figure 3.**
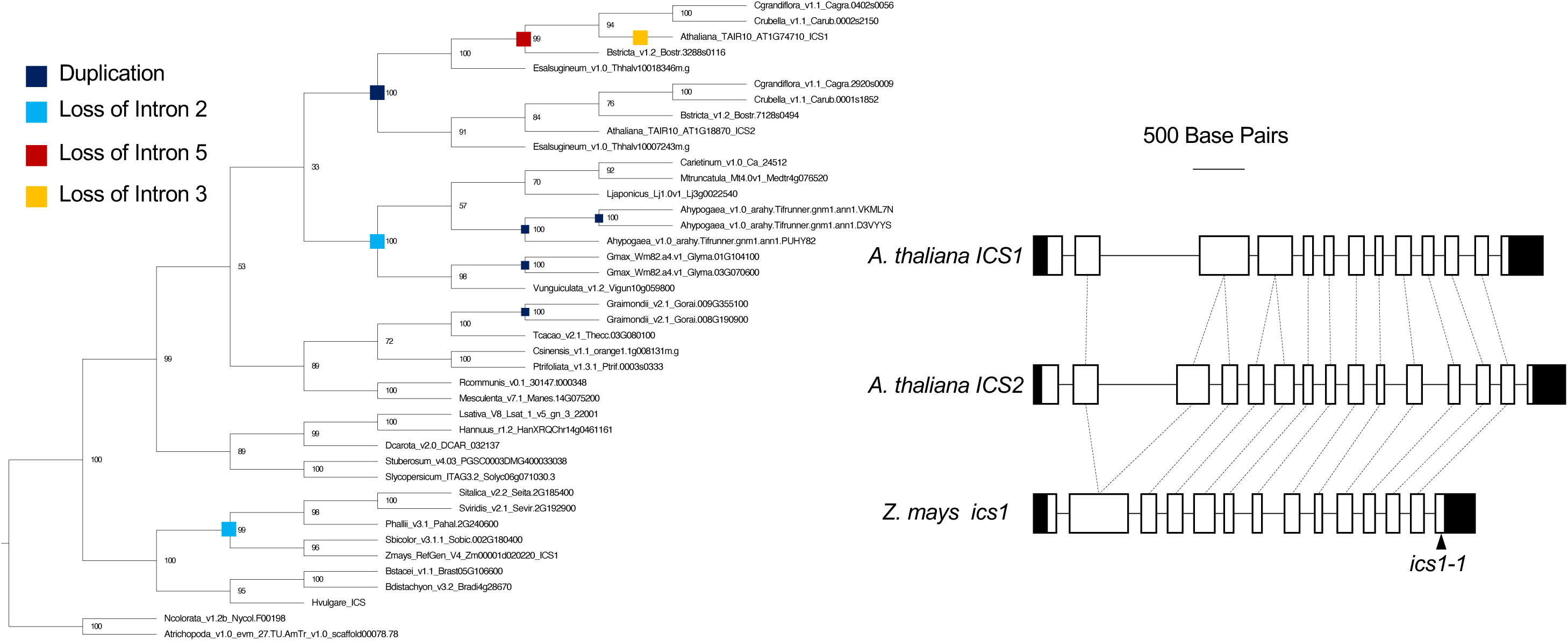
Phylogeny and exon/intron structures of angiosperm isochorismate synthase genes. All gene names are from Phytozome v13 except for *Hordeum vulgare.* Bootstrap values are out of 100 iterations. Colored squares represent points of gene duplication or loss of introns 2, 3, or 5 as depicted by the exon intron diagrams (right) for *Arabidopsis thaliana ICS1* and *ICS2* and *Zea mays ics1*.. Exons are denoted by empty white boxes and introns by thin black lines. Shaded boxes represent the 5’- and 3’-UTR. Exons connected by dotted lines denote regions of high sequence similarity. Mutator insertion position for *ics1-1*, relative to the *Z. mays ics1* gene structure, is labeled by a black arrowhead.

### Zea mays ics1 is required for phylloquinone biosynthesis

To test the function of maize *ics1*, we obtained a mutant, *ics1-1* (mu1039765), that contained a *Mutator* transposable-element (*Mu*) insertion in exon 14 of *ics1* (Figures 2 and 3). This mutation was inherited as a single recessive trait (wildtype:*ics1-1* - 51:17 in an F2 family resulting from a self-pollinated *ics1-1/+* heterozygous individual, χ^2^= 0, df = 1, *p* = 1). The *ics1-1* mutants were visually indistinguishable from wild-type sibling plants at emergence and did not show a yellow or stunted growth phenotype until 12-14 days after sowing (Figure 4). The yellow and stunted phenotype of the maize *ics1-1* mutant (Figures 4 and 5) was similar to that described previously for loss-of-function *ics* mutants in Arabidopsis and barley (Garcion et al., 2008; Qin et al., 2019). No recombinants were detected between the yellow leaf phenotype and the *Mu* insertion in *ics1-1* (76 green homozygous wild-type or heterozygous plants: 22 yellow homozygous mutants). The coloration of *ics1-1* mutants was consistent as the plants aged; however, the stunted growth phenotype became more severe (Figure 5A). The penetrance of the *ics1-1* mutant was extremely variable. Most *ics1-1* mutants died prior to reproduction or were sterile but rare individuals survived to maturity and produced viable pollen. To determine the consequence of *ics1* disruption on phylloquinone biosynthesis, we measured the amount of phylloquinone in leaves of wild-type and *ics1-1* mutant plants. The *ics1-1* mutants accumulated approximately 10% of wild-type phylloquinone levels (Figure 6A).

**Figure 4.**
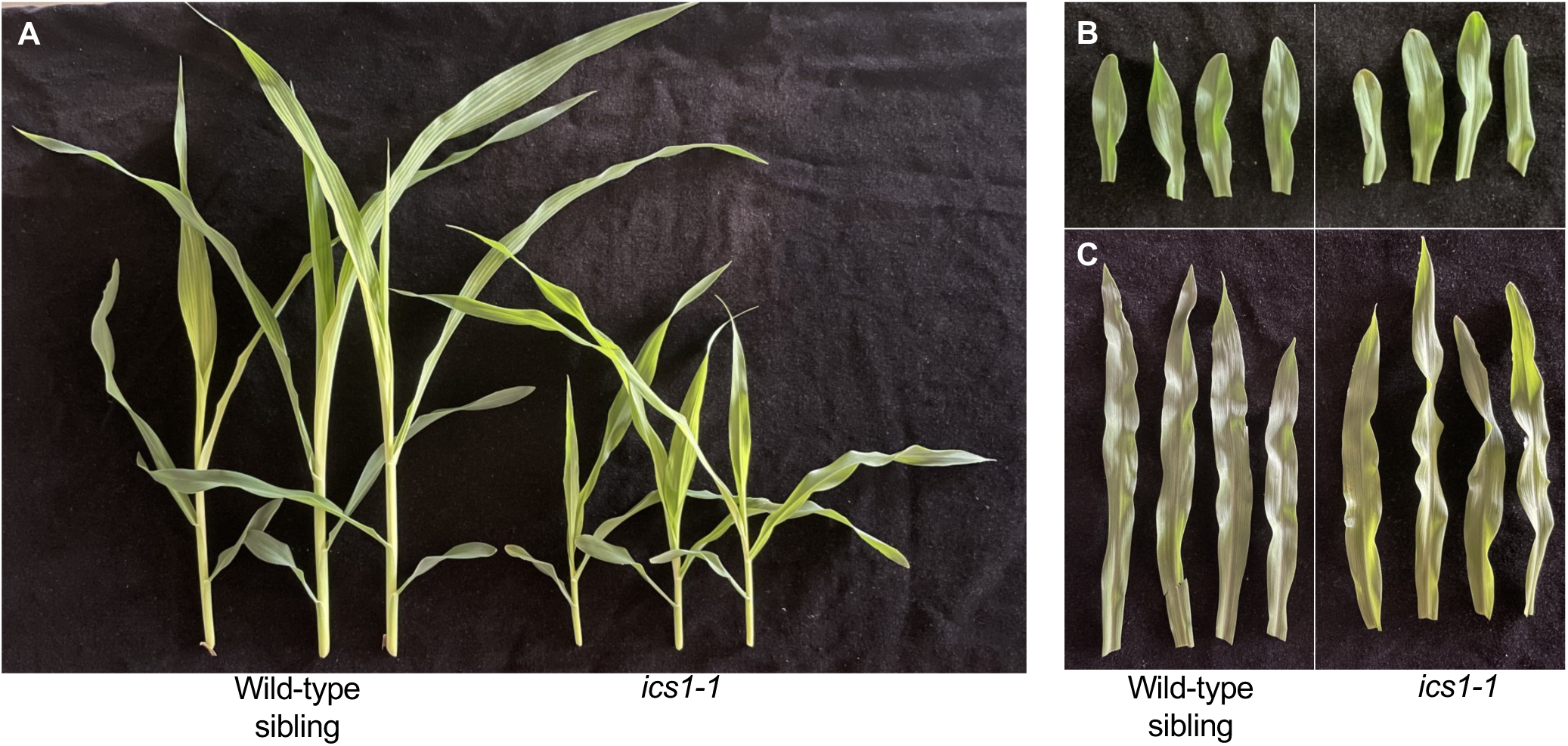
Phenotypic severity of *ics1-1* mutants. Representative (A) whole seedlings, (B) first leaves, and (C) second leaves from 14-day old *ics1-1* mutants and wild-type siblings.

**Figure 5.**
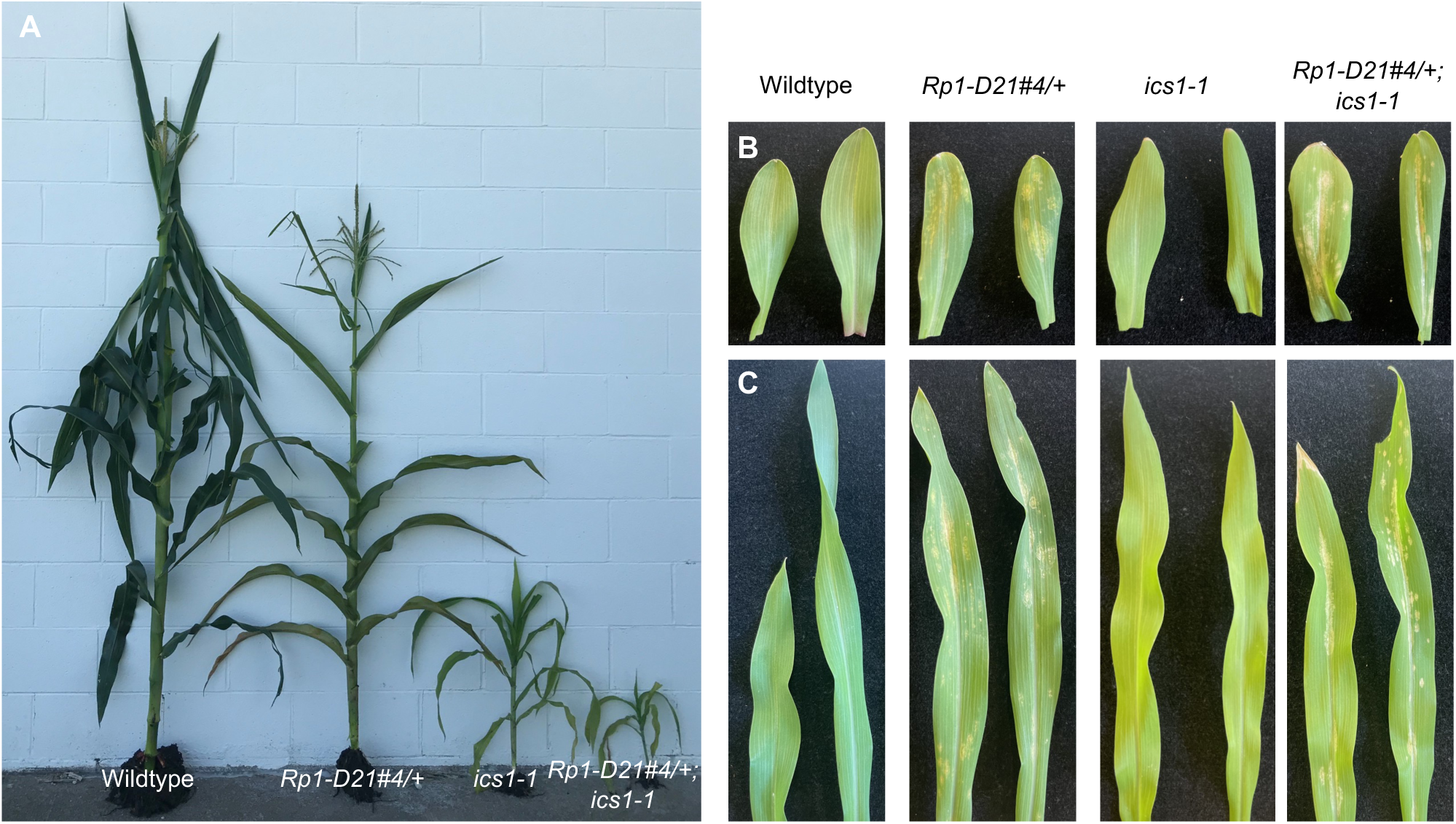
The *ics1-1* mutant is stunted and pale, and the *Rp1-D21#4/+; ics1-1* double mutant has a similar lesion severity as the *Rp1-D21#4/+* mutant. (A) Field grown wild-type, *Rp1-D21#4/+*, *ics1-1*, and *Rp1-D21#4/+; ics1-1* plants ∼50 days after planting. Representative (B) first and (C) second leaves from wildtype, *Rp1-D21#4/+* mutants, *ics1-1* mutants, and *Rp1-D21#4/+;ics1-1* double mutants grown under greenhouse conditions 16-days after sowing.

**Figure 6.**
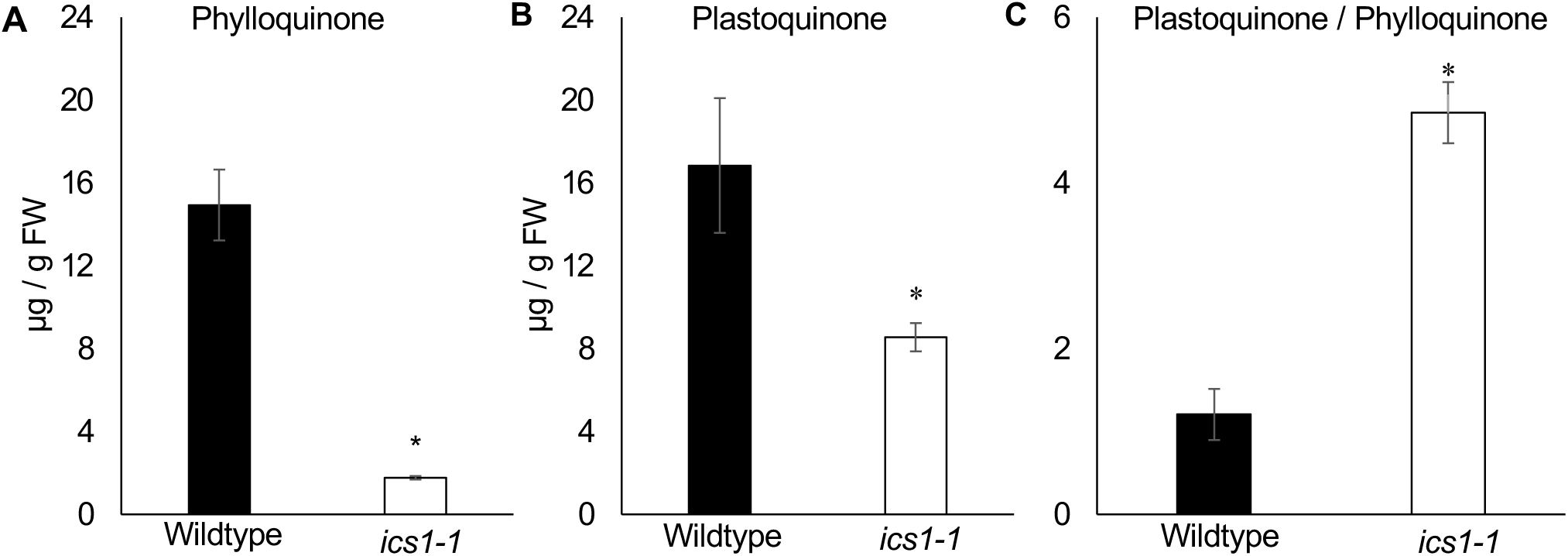
Phylloquinone and plastoquinone accumulation is reduced in the *ics1-1* mutant. (A) Phylloquinone and (B) plastoquinone amounts and (C) and the relative ratio of plastoquinone/phylloquinone in wild-type and *ics1-1* siblings. Data presented are mean ± S.E. (n = 4). Stars above bars represent statistically different measurements between wildtype and *ics1-1* (Student’s T-test, *p* < 0.05).

### ics1-1 is deficient in electron transport and has decreased amounts of Photosystem I

We next evaluated multiple photosynthetic parameters for *ics1-1* mutant and wild-type siblings to see if the loss of phylloquinone in *ics1-1* mutants disrupted photosynthesis. Compared to wildtype, *ics1-1* mutants had increased non-photochemical quenching (NPQ; Figure 7A), indicating that *ics1-1* mutants cannot utilize light as efficiently as wild-type plants. In addition, the *ics1-1* mutants had decreased operating and maximum quantum efficiencies of PSII (ΦPSII and F_v_/F_m_; Figures 7B and 7D), and linear electron transport rate (ETR; Figure 7C) compared to wild-type plants, demonstrating lower photosynthetic efficiency in *ics1-1* mutants. The oxidation state of the plastoquinone (PQ) pool was also much lower in *ics1-1* mutants than in wild-type controls (Figure 7F).

**Figure 7.**
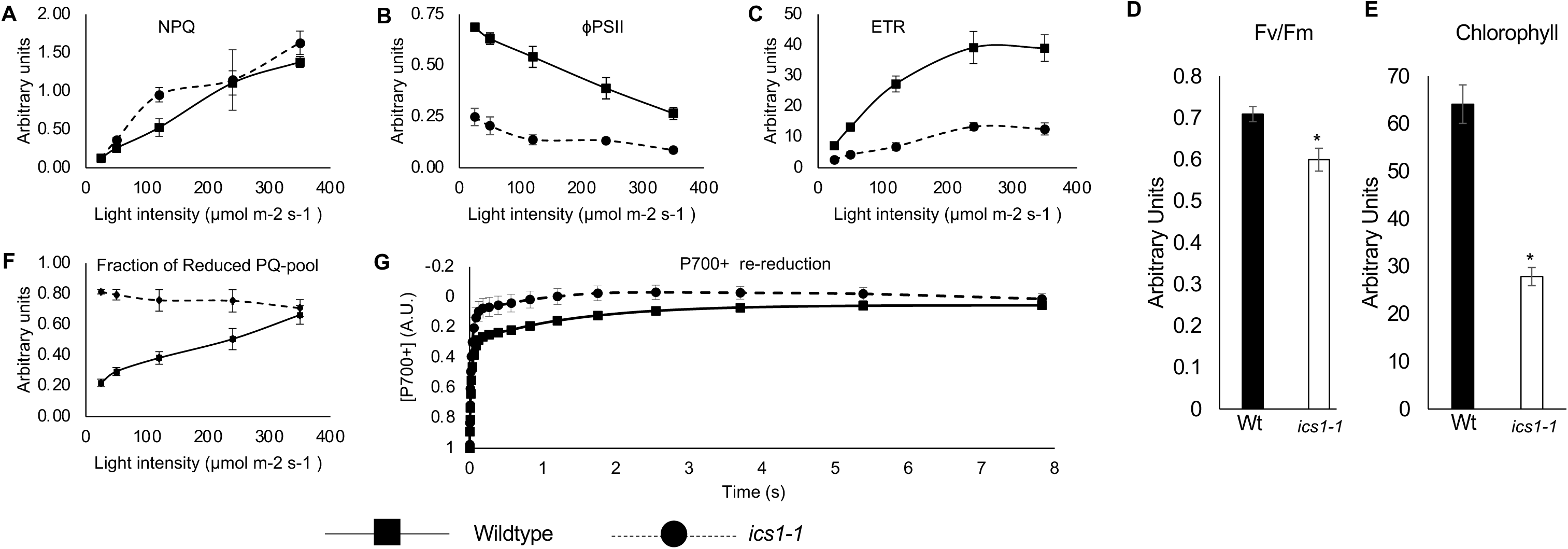
*ics1-1* mutants have decreased photosynthetic efficiency and impaired electron transport. (A) Non-photochemical quenching (NPQ), (B) PSII efficiency (ΦPSII), (C) electron transport rate (ETR), (D) Maximum efficiency of PSII photochemistry (Fv/Fm), (E) relative chlorophyll content, and (F) fraction of reduced plastoquinone (PQ) -pool in wild-type and *ics1-1* siblings. Data presented are means ± S.E. (n ≥ 5). Stars above bars indicate a statistically significant difference between wild-type and *ics1-1* samples (Student’s T-test, *p* < 0.05). (G) Re-reduction of P700^+^ complexes in wild-type and *ics1-1* siblings over time after transfer from light to dark. Data for each timepoint represents the mean ± S.E. (n > <u>5</u>).

Given phylloquinone’s role as an electron carrier in PSI, we evaluated how the loss of phylloquinone alters PSI electron transport in *ics1-1* mutants by measuring the rate of re-reduction of the reaction center P700^+^ chlorophyll in *ics1-1* mutants and wild-type plants (Figure 7G). Plastoquinone can occupy PSI in the absence of phylloquinone in both *Synechocystis sp.* PCC 6803 (Semenov et al., 2000) and *Chlamydomonas reinhardtii* (Lefebvre-Legendre et al., 2007). The replacement of phylloquinone with plastoquinone reduces the free energy gap to the F_x_ iron-sulfur cluster in PSI, resulting in a faster P700 dark re-reduction due to a backflow of electrons from plastoquinone to P700^+^ (Johnson et al., 2000; Shinkarev et al., 2002). The dark re-reduction of the P700^+^ complex was faster in *ics1-1* mutants than in wild-type controls, indicating that forward electron transport through PSI is altered in *ics1-1* mutants (Figure 7G). These findings are consistent with an electron carrier with a more positive midpoint potential, such as plastoquinone, replacing phylloquinone in PSI in *ics1-1* mutants. The amount of plastoquinone in leaves of *ics1-1* mutants is half that of wild-type samples (Figure 6B); however, the 2-fold reduction of plastoquinone and the 8.4-fold reduction of phylloquinone in *ics1-1* mutants compared to wild-type samples results in the relative ratio of plastoquinone to phylloquinone changing from 1:1.1 in wild-type plants to 4.8:1 in *ics1-1* mutants (Figure 6C). Therefore, despite a decrease in plastoquinone levels in *ics1-1* mutants, plastoquinone is 4.8-times more abundant than phylloquinone in these mutants which may facilitate plastoquinone replacing phylloquinone in PSI. Like the maize *ics1-1* mutant, the *Arabidopsis ics1; ics2* double mutants also reduced plastoquinone accumulation compared to wildtype (Gross et al., 2006), which indicates that plastoquinone biosynthesis has a yet-to-be characterized requirement for ICS.

Finally, we evaluated how disruption of *ics1* alters the stoichiometric ratios of PSII and PSI. On a per chlorophyll basis, *ics1-1* mutants have about 2-times more PSII (Figure 8A) and about 2.7-times less PSI (Figure 8B) than wild-type plants. This results in a shift in the ratio of PSII:PSI from 1:2 in wild-type plants to 2.7:1 in *ics1-1* mutants (Figure 8C). Since chlorophyll levels in *ics1-1* mutants are about half the level in wild-type plants (Figure 7E),the total amount of PSII in *ics1-1* mutants and wild-type plants is similar but the amount of PSI in *ics1-1* mutants is approximately 5.3-times less than wild type. This is similar to the observations and interpretation of phylloquinone loss mutants of *Chlamydomonas reinhardtii* (Sakuragi et al., 2002) Thus, the altered stoichiometry between PSII and PSI in *ics1-1* mutants is driven by the loss of PSI.

**Figure 8.**
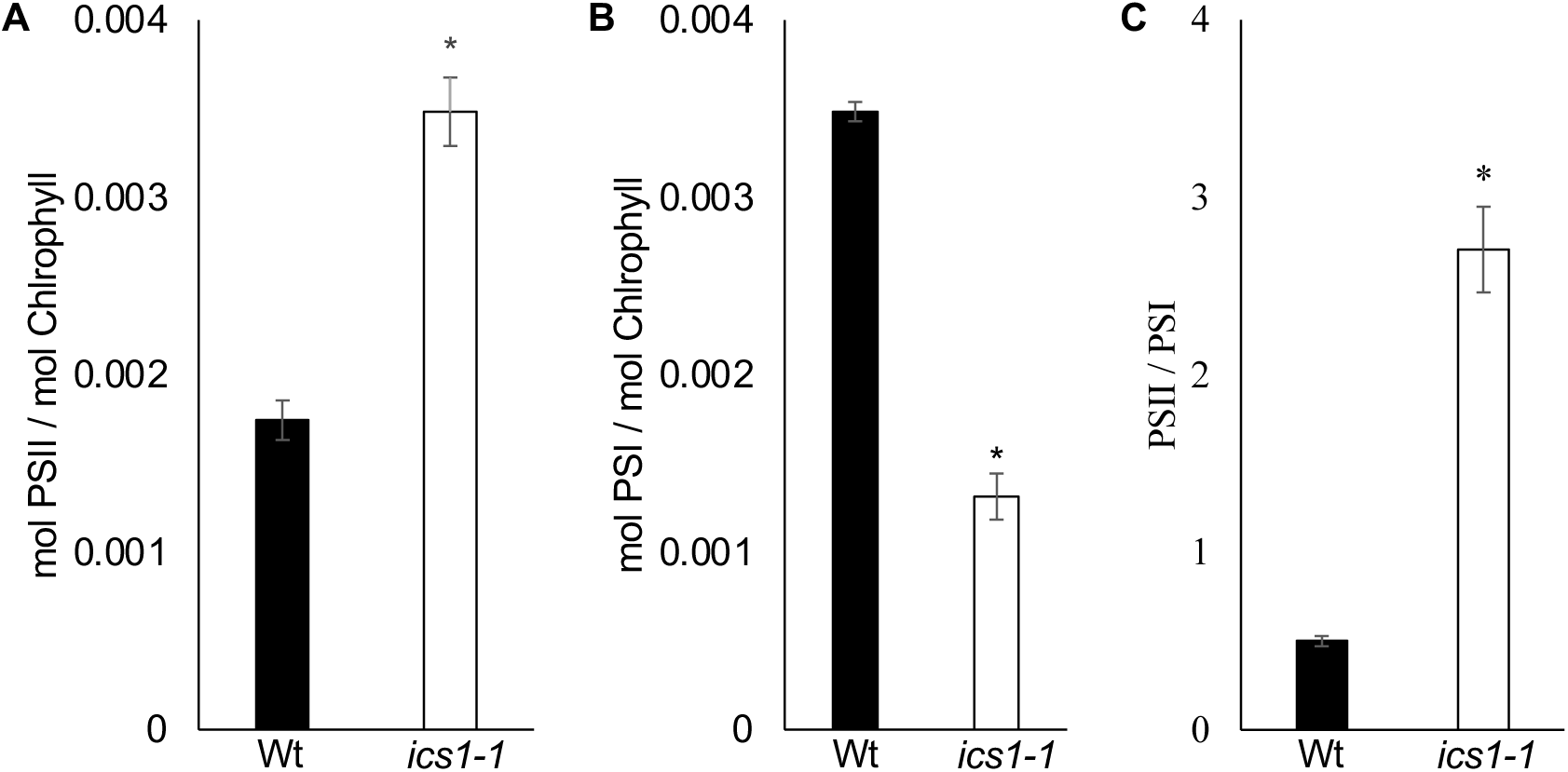
The stoichiometric ratio of PSII to PSI is altered *ics1-1* mutants. (A) Amount of photosystem II and (B) photosystem I relative to chlorophyll and (C) the relative ratios of photosystem II and I in Wt and *ics1-1* siblings. Data presented are means ± S.E. (n = 4). Stars above bars represent statistically different measurements between wild-type and *ics1-1* siblings (Student’s T-test, *p* < 0.05).

### ics1 is required for hypersensitive response-induced SA biosynthesis in maize

Disruption of *ICS* in plants blocks the biosynthesis of phylloquinone and SA (Figure 6A; Wildermuth et al., 2001; Gross et al., 2006; Garcion et al., 2008; Qin et al., 2019). In addition, loss of ICS activity indirectly leads to changes in the levels of other metabolites, such as lower levels of plastoquinone (Figure 6B; Gross et al., 2006) and chlorophyll (Figure 7E). To investigate the metabolic consequences of disrupting *ics1* in maize leaves, we performed a liquid-chromatography/mass-spectrometry (LC/MS) untargeted analysis of methanol-soluble metabolites separated by reverse phase chromatography. This analysis efficiently detects SA and multiple SA-derivatives while also enabling the analysis of thousands of additional metabolic features. In addition to the *ics1-1* mutant, we also included the *Rp1-D21#4/+* mutant in our metabolite analysis. *Rp1-D21#4* encodes a weak intragenic suppressor allele of *Rp1-D21* derived by ethyl methanesulfonate (EMS) mutagenesis that still displays constitutively active HR, but has a less severe phenotype and greater fertility than *Rp1-D21* (Wang and Balint-Kurti, 2015; Karre et al., 2021). Also included in this metabolite analysis were *Rp1-D21#4/+; ics1-1* double mutants, which permitted evaluation of both basal and HR-responsive SA biosynthesis and analysis of how other HR-responsive metabolites are affected by disruption of *ics1*.

The material used for this untargeted metabolite analysis was from the sibling progeny of a cross of +/+; *ics1-1/+* with *Rp1-D21#4/+*; *ics1-1/+* that segregated phenotypically 3:3:1:1 for wildtype, *Rp1-D21#4*, *ics1-1*, and *Rp1-D21#4; ics1-1* (58:54:20:10, χ^2^= 4.10, df = 3, *p* = 0.25). The untargeted metabolomics analysis of this tissue detected 7,094 mass features that were reproducibly detectable in all replicates (four for wild type and *Rp1-D21#4/+*, three for *ics1-1*, and five for *Rp1-D21#4/+; ics1-1*) of at least one of the four genotypes. Peak areas for all features were extracted using XCMS (Smith et al., 2006), and the average peak area of each feature was calculated for each of the four genotypes and compared for compounds of interest. SA was identified in our analysis by comparison to an authentic standard. Both the SA parent ion and spontaneous decarboxylation fragment were detected by MS with the decarboxylated fragment being the more abundant feature (M137.024_T538 and M93.034_T538 in Supplemental Data Set S1). Consistent with earlier reports of *Rp1-D21/+* mutants (Ge et al., 2021), SA accumulation was 4.5 times greater in *Rp1-D21#4/+* mutants compared to wild-type samples (Figure 9A). SA accumulation in *ics1-1* mutants and *Rp1-D21#4/+; ics1-1* double mutants was not significantly different from accumulation in wild-type samples indicating that *ics1-1* was epistatic to *Rp1-D21#4* for induced SA accumulation. The requirement for ICS1 was also observed for SA catabolites. SA-glucoside (Figure 9B, M299.07_T317), 2,5-DHBA glucoside (Figure 9C M315.072_T186), and 2,3-DHBA glucoside (Figure 9D, M315.072_T244; See methods for means of compound identification) are all compounds synthesized from SA in plants (Figure 1). All three accumulated in *Rp1-D21#4/+* mutants and were not significantly different between the *Rp1-D21#4/+; ics1-1* double mutants and wild-type controls. The two DHBA glucosides were also reduced in *ics1-1* mutants as compared to wild-type controls. Taken together, these findings demonstrate that *ics1* is required for *Rp1-D21#4*-induced SA accumulation.

**Figure 9.**
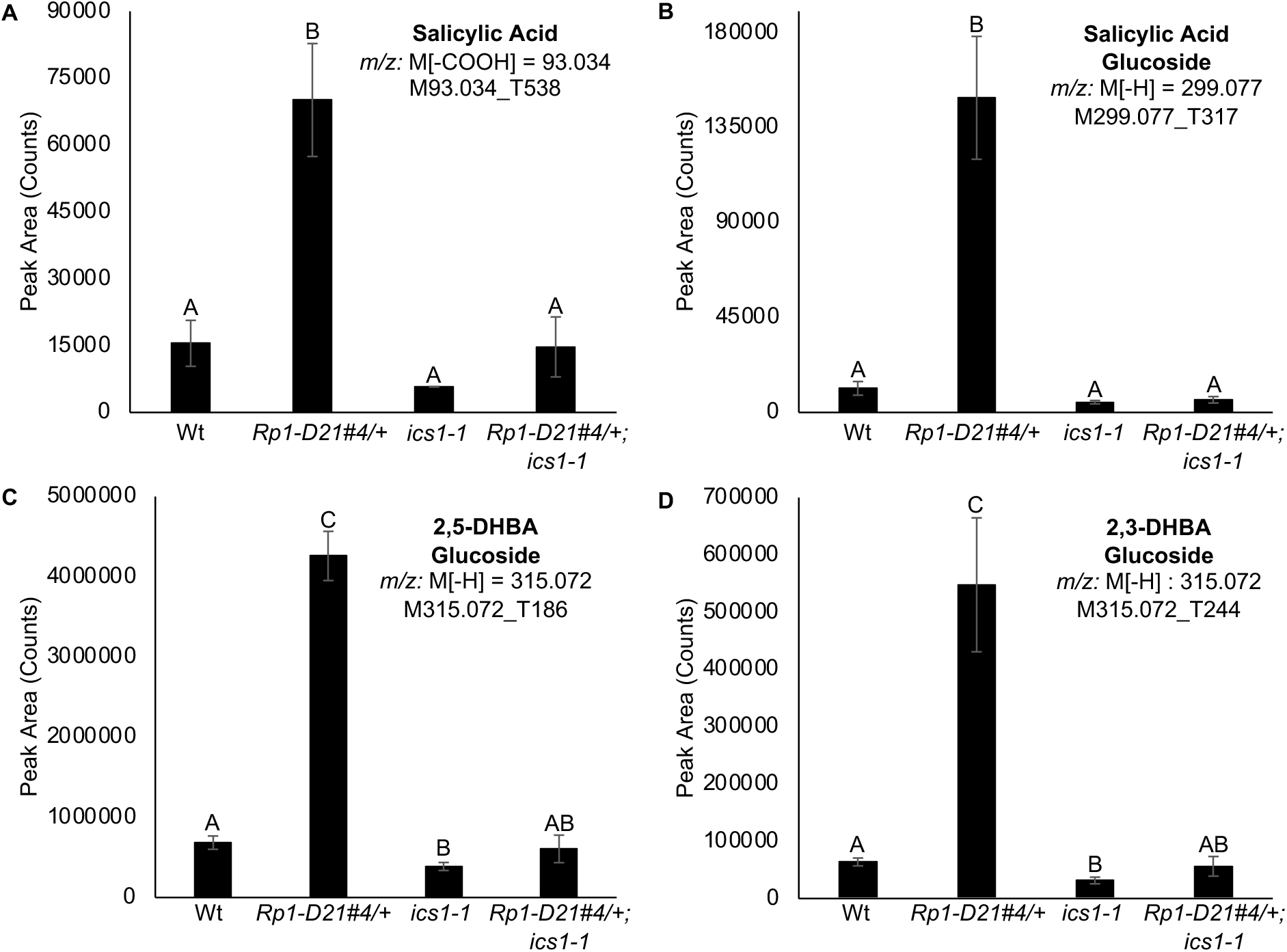
*ics1* is required for *Rp1-D21#4*-induced SA accumulation. Integrated peak areas of (A) salicylic acid, (B) salicylic acid glucoside, (C) 2,5-DHBA glucoside, and (D) 2,3-DHBA glucoside from leaves of wildtype (Wt), *Rp1-D21#4/+, ics1-1*, and *Rp1-D21#4/+; ics1-1*. Metabolites were analyzed in negative ionization mode. For salicylic acid, the integrated values are of the major decarboxylation fragment M[-COOH]. For salicylic acid glucoside, 2,5-DHBA glucoside, and 2,3-DHBA glucoside the values are integrations of the parent ion. Data presented are the mean values ± S.E. (n ≥ 3). Letters above bars represent statistically different measurements (paired sample Student’s T-test, *p* < 0.05).

Previously, SA accumulation and HR severity were measured in *Rp1-D21/+* mutants crossed to different genetic backgrounds (Ge et al., 2021). We calculated the correlation between SA accumulation and lesion severity in these *Rp1-D21/+* mutants. Lesion severity was positively correlated with both free SA (*r* = 0.72, *p* = 0.01) and total SA (*r* = 0.92, *p* = 4.5E-5) levels (Supplemental Figure S1). The suppression of SA accumulation during *Rp1-D21#4*-induced HR by *ics1-1* allowed us to test whether SA accumulation was necessary for lesion formation during HR in maize. Lesion formation in *Rp1-D21#4/+; ics1-1* double mutants was visually indistinguishable from *Rp1-D21#4/+* single mutants (Figures 5B and 5C) demonstrating that SA accumulation was a response to lesion formation and increased accumulation of SA was not required for lesion formation during HR in maize.

The biosynthetic steps that comprise an ICS-dependent pathway to SA were recently elucidated in Arabidopsis (Rekhter et al., 2019; Torrens-Spence et al., 2019). Isochorismate is first conjugated with glutamate by PBS3 to form the intermediate isochorismoyl-glutamate which spontaneously fragments or is enzymatically cleaved by EPS1 to produce SA (Figure 1). Isochorismoyl-glutamate was readily detectable in 50% methanol extracts from Arabidopsis at an *m/z* [M-H] = 354.083 in negative ionization mode (Rekhter et al., 2019; Torrens-Spence et al., 2019). Despite analyzing mutants that accumulated SA using similar extraction and chromatography as the Arabidopsis studies, we did not detect any mass features that could be isochorismoyl-glutamate in any of our maize genotypes (Supplemental Data Sets 1, 3, 4, 6). It is possible that this pathway exists in maize but the intermediate does not accumulate or that a different amino acid conjugation is used. We calculated the expected *m/z* values for isochorismate conjugated with the other 19 amino acids and searched the untargeted mass feature list for any mass matches. Of all the matches, only one mass feature, with the predicted mass of a conjugate with threonine (M326.088_T171 in Supplemental Data set S1), increased in accumulation in *Rp1-D21#4/+* mutants. We used linear modeling to test the interaction between *Rp1-D21#4* and *ics1-1* to determine if accumulation of this feature was dependent on *ics1-1*. The M326.088_T171 feature accumulation in *Rp1-D21#4* was partially dependent on *ics1-1* and *Rp1-D21#4 ics1-1* double mutants accumulated less of this feature than the *Rp1-D21#4* single mutant (Supplemental Data set S1 and summarized in Supplemental Table S1). The failure to detect isochorismoyl-glutamate, especially in *Rp1-D21#4* mutant plants that accumulate SA, suggests that this pathway is not responsible for induced SA accumulation in maize.

### SA derivatives are synthesized from phenylalanine in maize

Genetic evidence suggests that the PAL-dependent SA biosynthetic pathway may contribute as much as 50% of total SA biosynthesis in maize (Yuan et al., 2019); however, this has never been demonstrated biochemically. To test if SA can be synthesized from phenylalanine in maize, we performed stable isotopic labeling of *Rp1-D21/+* mutants and wild-type controls with phenylalanine or ^13^C-ring-labeled phenylalanine via roots. Soluble metabolites were extracted from developing shoots and analyzed using the same untargeted metabolomics analysis described above. Compounds synthesized from exogenously supplied ^13^C ring-labeled phenylalanine will co-elute with their unlabeled counterparts, but differ in *m/z* by 6.020 amu. Pools of the SA derivatives, SA glucoside (M299.077_T326 in supplemental Data Set S2), 2,5-DHBA glucoside (M315.072_T190), and 2,3-DHBA glucoside (M315.072_T295) were all labeled by ^13^C-phenylalanine by 1-2%. The labeled forms of these compounds were detected in plants fed ^13^C-ring-labeled phenylalanine and not in any plants fed non-labeled phenylalanine (Figure 10, Supplemental Figure S2). Although labeled conjugated SA was detected, labeled free SA was not detected as either the parent ion (Supplemental Figure S3) or decarboxylation fragment (Supplemental Figure S4) in any of the three *Rp1-D21*/+ replicates fed ^13^C-ring labeled phenylalanine. Taken together, the presence of labeled SA derivatives from fed ^13^C-ring-labeled phenylalanine suggests that SA can be produced from phenylalanine in maize.

**Figure 10.**
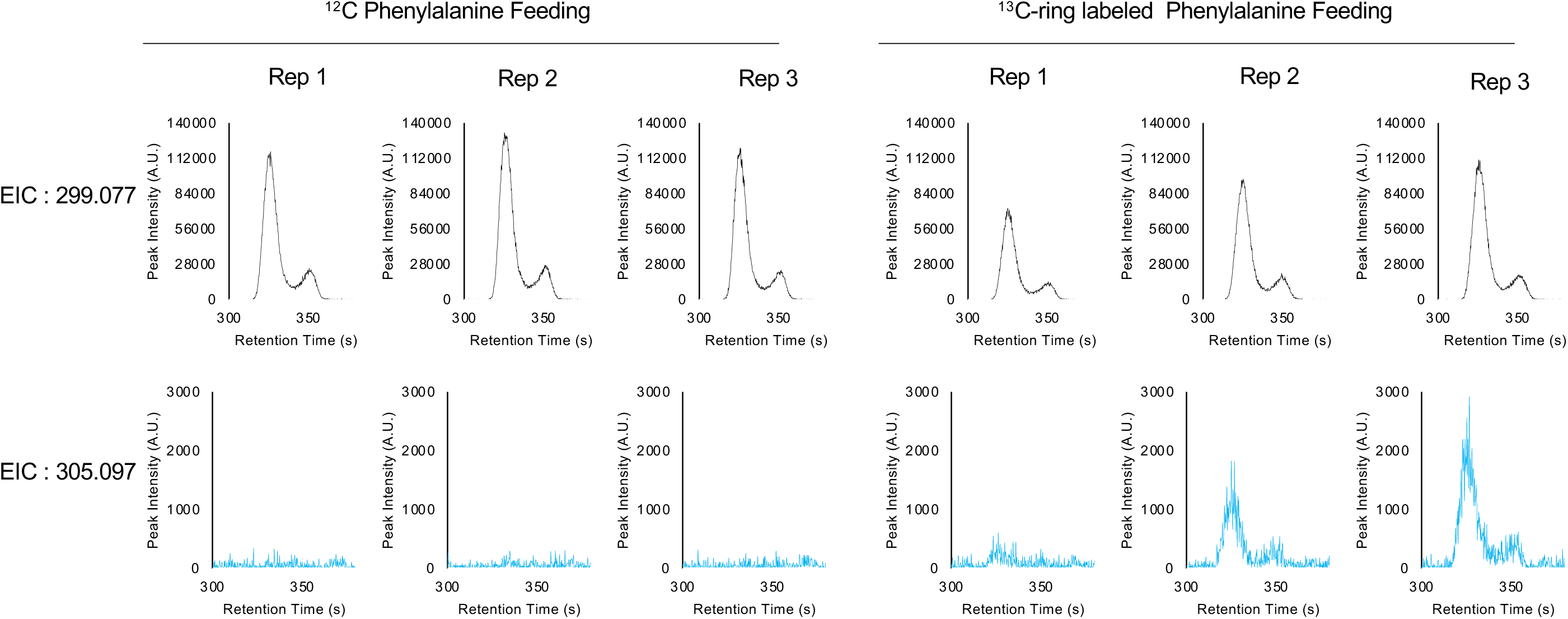
Salicylic acid glucoside can be synthesized from phenylalanine in maize. Extracted ion chromatograms (EIC) for (top) SA-glucoside (*m/z*: [-H] = 299.077) and (bottom) ^13^C-ring labeled SA-glucoside (*m/z*: [-H] = 305.097) for *Rp1-D21-ref* mutants fed (left) non-labeled phenylalanine or (right) ^13^C-ring labeled phenylalanine.

### Disruption of ics1 suppressed PAL activity, increased tryptophan and phenylalanine accumulation, and decreased accumulation of phenylalanine-derived compounds

To evaluate if disrupting *ics1* affects chorismate-derived metabolism, we compared the accumulation of L-tryptophan and L-phenylalanine, two products derived from chorismate (Figure 1), in wildtype, *Rp1-D21#4/+* mutants, *ics1-1* mutants, and *Rp1-D21#4/+; ics1-1* double mutants using data generated from our untargeted metabolomics data. Phenylalanine (M164.072_103 in Supplemental Data Set S1) and tryptophan (M203.038_T192) were identified by comparison to an authentic standard. Both amino acids accumulated to higher levels in *ics1-1* mutants compared to wild type (Figures 11A and 11B). In addition, *Rp1-D21#4/+; ics1-1* double mutants accumulated significantly higher levels of each amino acid compared to *ics1-1* mutants (Figures 11A and 11B). Thus, disrupting *ics1* resulted in an increased accumulation of chorismate-derived amino acids (Figure 1) consistent with increased substrate availability.

**Figure 11:**
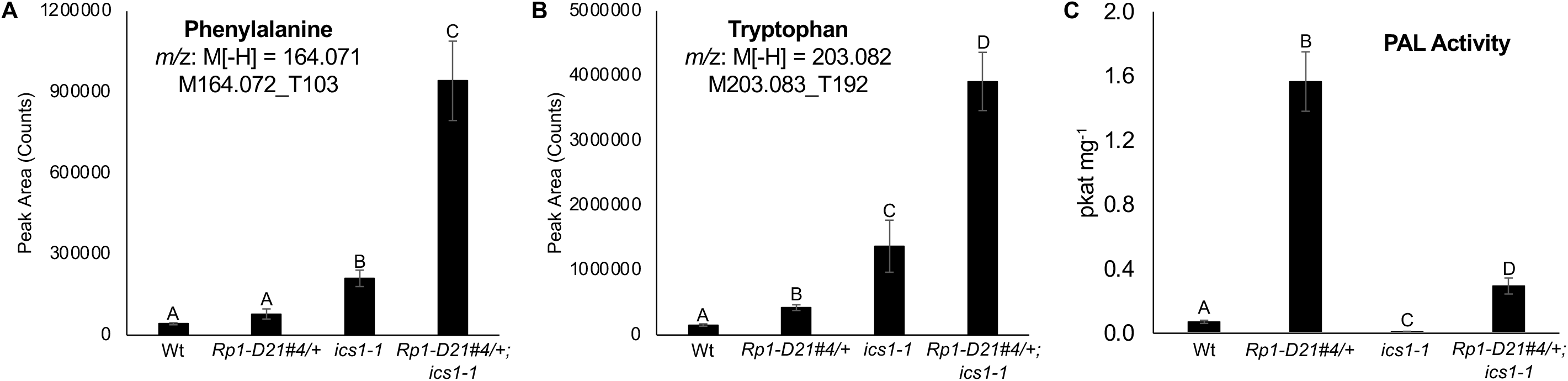
The *ics1-1* mutant has decreased PAL activity coinciding with increased accumulation of phenylalanine and tryptophan. (A) Phenylalanine accumulation, (B) tryptophan accumulation, and (C) PAL activity in wildtype (Wt), *Rp1-D21#4/+, ics1-1*, and *Rp1-D21#4/+; ics1-1* samples. Metabolites were analyzed in negative ionization mode and the integrated values are of the major parent ion. Data presented are the mean values ± S.E. (n ≥ 3). Letters above bars represent statistically different measurements (paired sample Student’s T-tests, *p* < 0.05).

Evidence of regulatory cross talk between the ICS-dependent and PAL-dependent SA biosynthetic pathways was previously observed in Arabidopsis. The combined decrease in SA accumulation of Arabidopsis *pal1/pal2/pal3/pal4* quadruple mutants and *ics1* mutants relative to wild-type samples exceeds 100% (Huang et al., 2010). Since disruption of ICS and PAL pathways each led to a greater than 50% reduction in SA accumulation in Arabidopsis, we propose that the ICS1 and PAL pathways are co-regulated. If this co-regulation is conserved in maize, then the requirement of *ics1* for SA accumulation in *Rp1-D21#4* might result from both a disruption of ICS-dependent SA biosynthesis and an indirect reduction of the PAL-dependent SA biosynthetic pathway.

To determine if disrupting *ics1* affects phenylalanine-derived metabolism, we compared the accumulation of phenylalanine-derived compounds in *Rp1-D21#4/+*, *ics1-1*, and the *Rp1-D21#4/+; ics1-1* double mutant using our untargeted metabolomics data. Features derived from phenylalanine were identified by matching the masses and retention times from our ^13^C-phenylalanine feeding experiment to our 7,094-feature list from our untargeted analysis. This identified 124 phenylalanine-derived features in our untargeted metabolite profiling. Three of these features corresponded to phenylalanine or fragments of it (M164.072_T116, M164.072_T103, and M147.045_T103 in Supplemental Data Set S1) and were not considered further in this analysis. To evaluate how the remaining 121 features were affected by genotype, we performed a linear model analysis that considered how *Rp1-D21#4*, *ics1-1*, and the interaction of *Rp1-D21#4* and *ics1-1* affect metabolite accumulation. The coefficients for *Rp1-D21#4* and *ics1-1* indicate the direction (positive: higher accumulating; negative: lower accumulating) and magnitude of change in mass feature accumulation in each genotype relative to the wild-type value. The coefficient for the *Rp1-D21#4* by *ics1-1* interaction indicates the direction and magnitude of how the accumulation of a mass feature deviates from the predicted additive effect of *Rp1-D21#4* and *ics1-1* in the *Rp1-D21#4/+; ics1-1* double mutant samples. For features where *Rp1-D21#4* and *ics1-1* affected accumulation in the same direction, we also considered in which genotype the magnitude of the affect was larger. Features were grouped by coefficient significance and the direction of the coefficients for *Rp1-D21#4*, *ics1-1*, and their interaction for all 121 features are presented in Table 1. Despite *ics1-1* mutants accumulating 5.1 times wild-type phenylalanine levels (Figure 11a), only 7 of the 121 features accumulated to significantly higher levels in *ics1-1* mutants compared to wild-type controls, whereas 28 features accumulated to significantly lower levels. Of the 121 features, 48 accumulated to significantly higher levels in *Rp1-D21#4/+* mutants than in wild-type controls. These findings are consistent with a previous RNA-seq analysis, which found that transcripts encoding steps in the phenylpropanoid pathway were increased in abundance in *Rp1-D21* mutants (Olukolu et al., 2014). Of these 48 features, 37 (77%) have a significant negative interaction effect, indicating *ics1-1* suppresses most *Rp1-D21#4*-induced phenylalanine-derived metabolite accumulation in *Rp1-D21#4/+; ics1-1* double mutants.

**Table 1:**
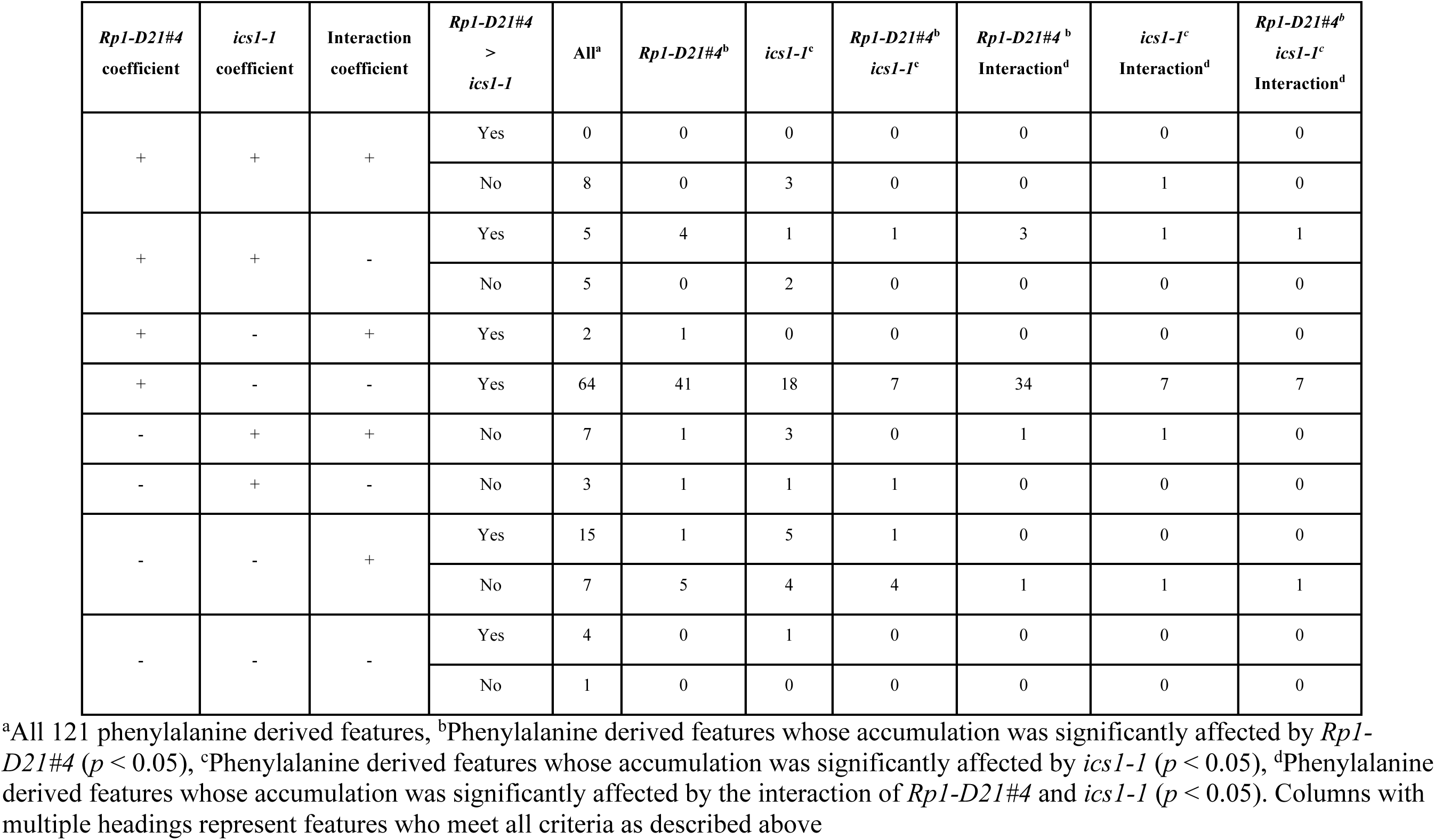
Disruption of *ics1* suppresses the accumulation of phenylalanine-derived metabolites

The increase in phenylalanine and decrease in phenylalanine-derived compounds in *ics1-1* mutants could be explained if maize PAL activity was reduced. To test if PAL activity is reduced in *ics1-1* mutants, we grew progeny segregating for wild type, *Rp1-D21#4/+*, *ics1-1*, and *Rp1-D21#4/+; ics1-1* and tested PAL activity in protein extracts from leaves of these four genotypes (Figure 11C). The *Rp1-D21#4/+* mutants had 22.2-fold higher PAL activity than wild-type siblings, consistent with the accumulation of many phenylalanine-derived compounds in *Rp1-D21#4/+* mutants. The *ics1-1* mutants had 8.2-fold less PAL activity than wild-type siblings. Consistent with the suppression of phenylalanine-derived metabolism in *ics1-1* mutants, PAL activity of the *Rp1-D21#4/+*; *ics1-1* double mutant was more than 5-fold less than that of *Rp1-D21#4/+* mutants. The reduction in PAL activity combined with the accumulation of phenylalanine and decrease in phenylalanine-derived compounds in the *ics1-1* mutant suggests that phenylalanine network metabolism is coordinately linked to isochorismate metabolism.

### Disruption of ics1 *suppresses* HR-responsive metabolism in Rp1-D21#4 mutants

To discern if *ics1-1* is epistatic to *Rp1-D21#4* for metabolites other than those derived from phenylalanine, we utilized the same linear model analysis and applied it to all 7,094 features and again assessed how *Rp1-D21#4*, *ics1-1*, and the interaction of *Rp1-D21#4* and *ics1-1* affect metabolite accumulation We used a false discovery rate (FDR) corrected (Benjamini and Hochberg, 1995) cutoff of *p* < 0.05 which retained 3,881 features whose accumulation was under genetic control. Features were grouped by coefficient significance and the direction of the coefficients for *Rp1-D21#4*, *ics1-1*, and their interaction for all 3,881 features are presented in Table 2.

**Table 2:**
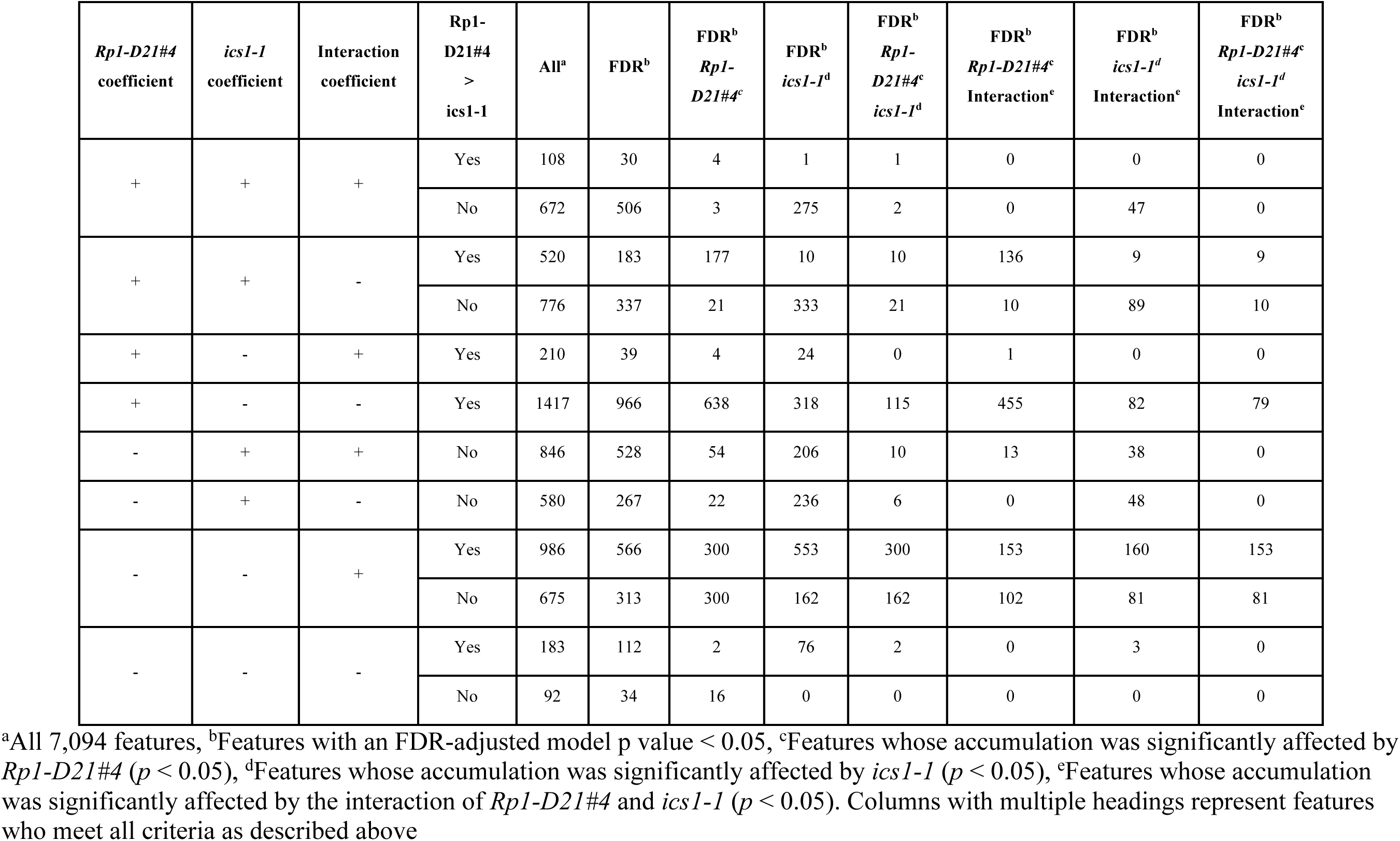
*Rp1-D21#4*-responsive metabolism is suppressed in *Rp1-D21#4 ;ics1-1* double mutants

| *Rp1-D21#4*<br>coefficient | *ics1-1*<br>coefficient | Interaction<br>coefficient | *Rp1-D21#4*<br>><br>*ics1-1* | All<sup>a</sup> | FDR<sup>b</sup> | FDR<sup>b</sup><br>*Rp1-D21#4*<sup>c</sup> | FDR<sup>b</sup><br>*ics1-1*<sup>d</sup> | FDR<sup>b</sup><br>*Rp1-D21#4*<br>*ics1-1*<sup>d</sup> | FDR<sup>b</sup><br>*Rp1-D21#4*<br>Interaction<sup>e</sup> | FDR<sup>b</sup><br>*ics1-1*<br>Interaction<sup>e</sup> | FDR<sup>b</sup><br>*Rp1-D21#4*<br>*ics1-1*<br>Interaction<sup>e</sup> |
| --- | --- | --- | --- | --- | --- | --- | --- | --- | --- | --- | --- |
| + | + | + | Yes | 108 | 30 | 4 | 1 | 1 | 0 | 0 | 0 |
|  |  |  | No | 672 | 506 | 3 | 275 | 2 | 0 | 47 | 0 |
| + | + | - | Yes | 520 | 183 | 177 | 10 | 10 | 136 | 9 | 9 |
|  |  |  | No | 776 | 337 | 21 | 333 | 21 | 10 | 89 | 10 |
| + | - | + | Yes | 210 | 39 | 4 | 24 | 0 | 1 | 0 | 0 |
| + | - | - | Yes | 1417 | 966 | 638 | 318 | 115 | 455 | 82 | 79 |
| - | + | + | No | 846 | 528 | 54 | 206 | 10 | 13 | 38 | 0 |
| - | + | - | No | 580 | 267 | 22 | 236 | 6 | 0 | 48 | 0 |
| - | - | + | Yes | 986 | 566 | 300 | 553 | 300 | 153 | 160 | 153 |
|  |  |  | No | 675 | 313 | 300 | 162 | 162 | 102 | 81 | 81 |
| - | - | - | Yes | 183 | 112 | 2 | 76 | 2 | 0 | 3 | 0 |
|  |  |  | No | 92 | 34 | 16 | 0 | 0 | 0 | 0 | 0 |

<sup>a</sup>All 7,094 features, <sup>b</sup>Features with an FDR-adjusted model p value < 0.05, <sup>c</sup>Features whose accumulation was significantly affected by *Rp1-D21#4* ( $p < 0.05$ ), <sup>d</sup>Features whose accumulation was significantly affected by *ics1-1* ( $p < 0.05$ ), <sup>e</sup>Features whose accumulation was significantly affected by the interaction of *Rp1-D21#4* and *ics1-1* ( $p < 0.05$ ). Columns with multiple headings represent features who meet all criteria as described above

There were 847 features that accumulated to significantly higher levels (positive coefficient; *p* < 0.05) in *Rp1-D21#4/+* mutants and 1,061 features that accumulated to significantly higher levels in *ics1-1* mutants. Only 34 features accumulated to significantly higher levels in both genotypes, indicating that *Rp1-D21#4* and *ics1-1* predominantly lead to changes in the accumulation of different metabolites. The metabolites that were significantly decreased in abundance in either genotype did not show this pattern. Of the 694 and 1,113 features that accumulated to significantly lower levels in *Rp1-D21#4/+* and *ics1-1* mutants, respectively, 464 features accumulated to significantly lower levels in both mutants. Thus, *Rp1-D21#4* and *ics1-1* induced the accumulation of distinct metabolite sets but tended to decrease the abundance of a similar set of metabolites.

We next evaluated the interaction effects between *Rp1-D21#4* and *ics1-1*. There were 79 features that had significant and opposite effects of *Rp1-D21#4* and i*cs1-1* and significant interaction effects. In all 79 instances, *Rp1-D21#4* increased, and *ics1-1* decreased feature accumulation (Table 2). For all 79 features, *ics1-1* was epistatic to *Rp1-D21#4,* and the directions of their interaction effects were always in the same direction as the main effects for *ics1-1* (Table 2).

The epistatic interactions between *ics1-1* to *Rp1-D21#4* were further explored by analyzing the directions of genetic effects for all mass features affected by *Rp1-D21#4*. Of the 847 features that accumulated to significantly higher levels in *Rp1-D21#4*, 601 features (71%) had a significant negative interaction coefficient and only 1 feature (0.1%) had a significant positive interaction coefficient (Table 2). Thus, *ics1* was required to accumulate *Rp1-D21#4*-induced metabolites. Since only 79 of these 601 features had a main effect for *ics1-1*, these findings indicate that disruption of *ics1* suppressed *Rp1-D21#4*-induced metabolism despite not detectably altering basal accumulation of these features.

Consistent with *ics1-1* acting epistatically to *Rp1-D21#4* for many metabolite features (Table 2), the overall metabolic consequences of *Rp1-D21#4/+; ics1-1* double mutants were very similar to those of *ics1-1* mutants and not similar to *Rp1-D21#4/+* mutants. We calculated the log_2_ fold-change accumulation of each of the 3,881 features (Supplemental Data Set S1) for each of the three genotypes in relation to the wild-type controls and performed a linear regression analysis. The correlation coefficient between *Rp1-D21#4/+; ics1-1* and *ics1-1* was 0.93 (*p* < 10^-200^) while the correlation coefficient between *Rp1-D21#4/+; ics1-1* and *Rp1-D21#4/+* was only 0.15 (*p* = 8×10^-22^).

### Foliar application of SA does not restore HR-responsive metabolism to Rp1-D21#4/+; ics1-1 double mutants

SA is a known disease signaling molecule in plants, and suppression of HR-responsive metabolic perturbations by *ics1-1* in *Rp1-D21#4/+; ics1-1* double mutants may be the result of a failure to accumulate SA. To test if increasing SA levels in *Rp1-D21#4/+; ics1-1* double mutants could restore *Rp1-D21#4*-responsive metabolism, we first determined the effects of SA application on maize metabolism. Seedlings of the inbred B73 were sprayed 10 d after sowing with 0.1 mM SA, 1 mM SA, 5 mM SA, or mock solution every 24 h for 3 d. 72 hours after the first treatment, we performed an untargeted metabolite analysis on leaf samples. Measurements of free SA (decarboxylated fragment at M93.035_T643 in Supplemental Data Set S3; Supplemental Figure S5A) likely included SA accumulated on the leaf surface from the spray treatment. To better estimate the uptake and utilization of SA, we examined the derivatives SA-glucoside (M299.078_T412; Supplemental Figure S5B) 2,5-DHBA glucoside (M315.073_T235; Supplemental Figure S5C), and 2,3-DHBA glucoside (M315.072_T325; Supplemental Figure S5D). Levels of these three SA-derivatives increased in a dose-dependent manner. The 1 mM and 5 mM treatments accumulated levels of these metabolites comparable to or higher than those observed in *Rp1-D21/+* mutants (Supplemental Figure S5). Leaves were damaged and browned on plants treated with 5 mM SA, but not 1 mM, (Supplemental Figure S6), indicating that a high concentration of SA was toxic. Thus, the 1 mM SA spray was used in subsequent treatments.

Families segregating for wildtype, *Rp1-D21#4/+*, *ics1-1*, and *Rp1-D21#4/+; ics1-1* were grown for 14 days and then sprayed with control solution or 1 mM SA once every 24 hours for 3 days. After the third day, we performed an untargeted metabolomic analysis of leaves. In this analysis, we detected 6,676 mass features that were present in all replicates of at least one genotype/treatment combination. Consistent with our prior SA treatments, the levels of SA (M93.035_T642 in Supplemental Data Set S4), SA glucoside (M299.078_T411), 2,5-DHBA glucoside (315.073_T248), and 2,3-DHBA glucoside (M315.072_T323) were higher in all four genotypes in treated compared to non-treated samples (Figure 12).

**Figure 12:**
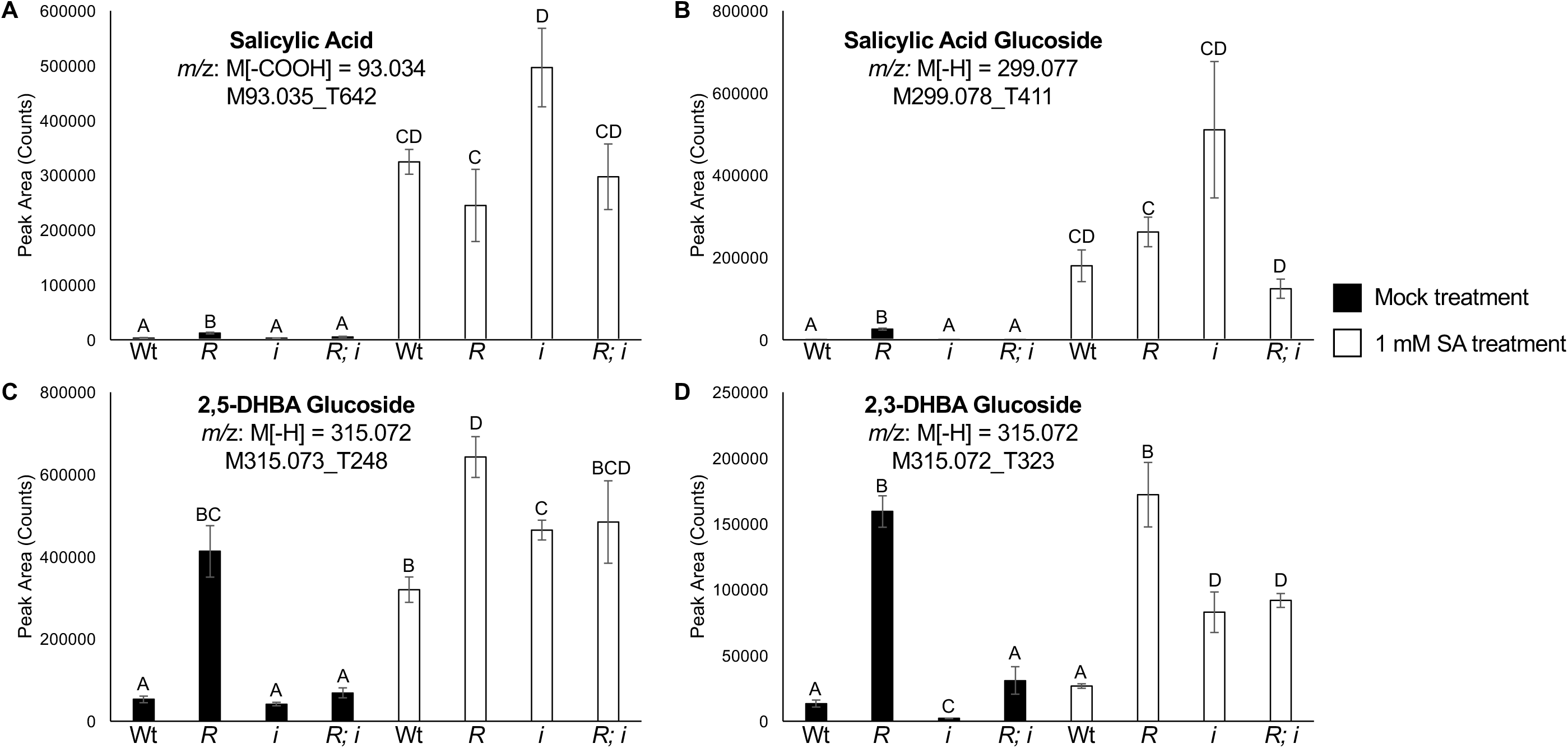
Foliar application of SA to maize leaves results in increased accumulation of SA and SA-derived compounds. Integrated peak areas of (A) salicylic acid, (B) salicylic acid glucoside, (C) 2,5-DHBA glucoside, and (D) 2,3-DHBA glucoside for wildtype (Wt), *Rp1-D21#4/+* (*R*), *ics1-1* (*i*) and *Rp1-D21#4/+; ics1-1* (*R;i*) plants treated with 0 mM, 0.1 mM, 1 mM, and 5 mM SA every 24 hours four three days. Metabolites were analyzed in negative ionization mode. For salicylic acid, the integrated values are of the major decarboxylation fragment M[-COOH]. For salicylic acid glucoside, 2,5-DHBA glucoside, and 2,3-DHBA glucoside the values are integrations of the parent ion. Data presented are the mean values ± S.E. (n = 3). Letters above bars represent statistically different measurements (paired sample Student’s T-test *p* < 0.05).

Metabolite features differentially affected by SA application were identified in these samples. Foliar application of 1 mM SA affected the levels of 308 features in wild-type samples (Student’s t-test, SA-treated vs. mock-treated, *p* < 0.05). Of these 308 features, 234 and 74 features showed increased and decreased accumulation respectively in wild-type plants after SA treatment. A greater number of metabolites, 505, were affected by 1 mM SA application to *ics1-1* mutants. Of these 505 features, 311 and 194 features showed increased and decreased accumulation respectively in *ics1-1* mutants after SA treatment. There were 74 features affected by SA treatment of both wild-type plants and *ics1-1* mutants. Of these 74 features, the SA-induced change in metabolite accumulation was in the same direction for 58 features and the opposite direction for 16 features (Chi-squared deviation from 1:1, *p* < 10^-7^). We next evaluated how the SA-responsive metabolites are affected by disruption of *ics1*. Of the 308 features affected by 1 mM SA supplementation in wild-type plants, the effect of *ics1-1* was in the same direction as the SA-treatment effect for 142 features and in the opposite direction for 166 features (Chi-squared deviation from 1:1, *p* > 0.05). These findings indicate that SA supplementation of wild-type plants did not predict the effect of disruption of *ics1*. Of the 505 features affected by 1 mM SA application to *ics1-1* mutants, the main effect of *ics1-1* was in the same direction as the SA-effect for 199 features and in the opposite direction for 306 features (Chi squared deviation from 1:1 p value< 10^-9^). Thus, the loss of *ics1-1* and supplementation of *ics1-1* mutants with SA had opposing effects for more metabolites than expected by chance, indicating SA supplementation was able to counteract the metabolic consequences of *ics1-1*.

If a lack of SA is the cause of the suppression of *Rp1-D21#4*-induced metabolite features in *Rp1-D21#4/+; ics1-1* double mutants, then features that accumulate in *Rp1-D21#4/+* and not in *Rp1-D21#4/+; ics1-1* double mutants should show increased accumulation in samples treated with SA. We again utilized a linear model to analyze the genetic effects of and interactions between *Rp1-D21#4/+* and *ics1-1* on metabolite features in the mock-treated samples (Supplemental Table S2). Of the 6,676 mass features detected, 1239 mass features accumulated to higher levels in *Rp1-D21#4/+,* and 901 (73%) of these were suppressed by the interaction of *ics1-1* and *Rp1-D21#4*. Of these 901 features, only 42 (5%) were significantly higher in *Rp1-D21#4/+; ics1-1* double mutants treated with 1 mM SA than mock-treated double mutants (Table 3). The 42 features included SA (decarboxylation fragment-M93.035_T642 in Supplemental Data Set S4) and masses consistent with SA derivatives including methyl salicylate (M151.04_T611), SA glucoside (Parent-M299.078_T411, aglucone fragment-M137.025_T411, decarboxylation fragment-M93.035_T409, SA-glucoside dimer-M599.162_T406), 2,3-DHBA glucoside (M315.072_T323), amino SA (Parent-M152.012_T257, decarboxylation fragment-M108.022_T257), and methyl amino SA (M166.027_T336). In addition to the putative amino SA and decarboxylation fragment, eight differentially accumulated metabolite features (Supplemental Table S4) co-elute from 254-258 seconds, suggesting they are either artifacts generated by the putative amino SA or derive from a co-eluting unknown compound. These features included a mass consistent with catechol (M109.029_T256), suggesting an additional unknown compound with a dihydroxy benzene moiety that is also derived from SA co-elutes at this time. Taken together, this experiment demonstrates that SA was not sufficient to restore the accumulation of most of the *Rp1-D21#4*-responsive metabolites that are suppressed in *Rp1-D21#4/+; ics1-1* double mutants. Of the features restored by SA treatment, many are likely derived directly from SA indicating that the addition of SA provided substrate for their synthesis and did not induce the accumulation of these metabolites through signaling.

**Table 3:**
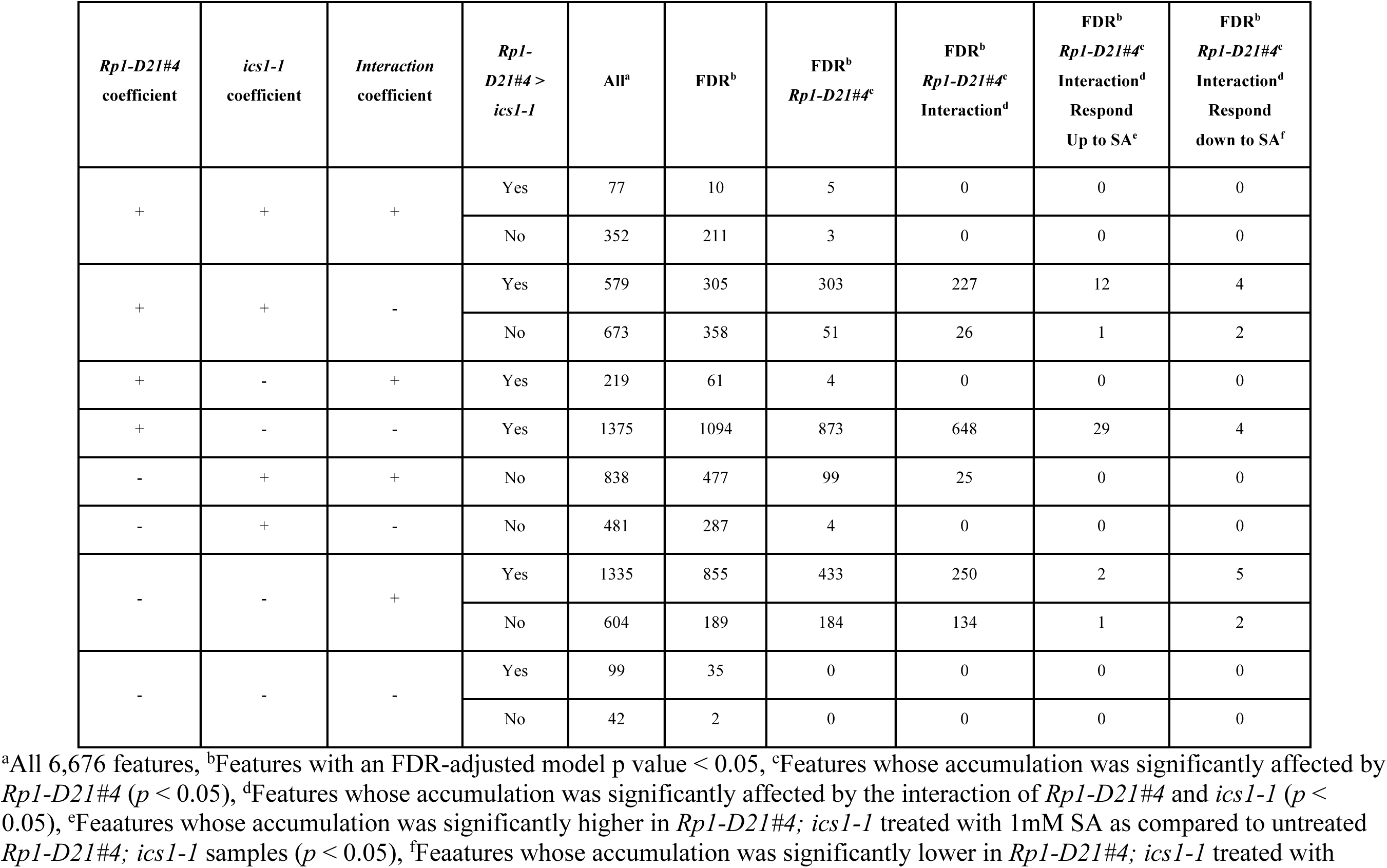
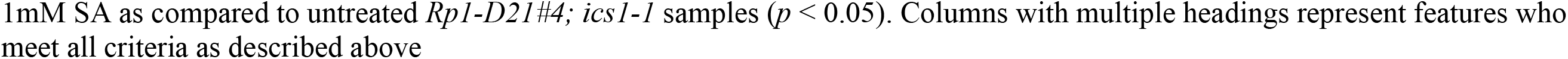
Most of the *Rp1-D21#4*-induced metabolites that are suppressed in *Rp1-D21#4; ics1-1* double mutants do not response to SA treatment.

### Loss of chlorophyll by Oy1-N1989 does not suppress Rp1-D21#4-induced metabolism

If the loss of photosynthetic capacity in *ics1-1*, and not a specific requirement for a downstream isochorismate-derived metabolite(s), is responsible for the suppression of *Rp1-D21#4*-induced metabolism, any mutant with impaired photosynthesis should also be epistatic to *Rp1-D21#4*. To test this, we used the *Oy1-N1989* mutant, a semi-dominant allele of subunit I of the Magnesium Chelatase enzyme in the chlorophyll pathway. Chlorophyll levels of *Oy1-N1989/+* mutants are only 10% of wild-type siblings early in development (Khangura et al., 2019). In addition, *Oy1-N1989* mutants have decreased CO_2_ assimilation and decreased maximum quantum yield of PSII (Fv/Fm; Khangura et al., 2020). We crossed *Oy1-N1989/+* mutants as a pollen parent onto *Rp1-D21#4/+* mutant ears to generate material segregating phenotypically for wild type*, Rp1-D21#4, Oy1-N1989,* and *Rp1-D21#4; Oy1-N1989* double mutants. The same untargeted metabolomic profiling described above was performed to determine what effect loss of chlorophyll and reduced photosynthesis had on metabolite accumulation in these leaf samples. This analysis detected 1,927 mass features that were reproducibly detectable in all replicates (n=4) in at least one of the four genotypes.

Like the *Rp1-D21#4/+* and *Rp1-D21#4/+; ics1-1* siblings, there were little to no distinguishable differences in the lesion phenotypes of *Rp1-D21#4/+* and *Rp1-D21#4/+; Oy1-N1989/+* siblings (Supplemental Figure S7). Unlike the *Rp1-D21#4/+; ics1-1* double mutants, the metabolite response of *Rp1-D21#4/+; Oy1-N1989/+* double mutants closely resembled that of *Rp1-D21#4/+* mutants. We calculated the log_2_ fold-change accumulation of all 1,927 features (Supplemental Data Set S5) for each genotype relative to the wild-type controls and performed a linear regression analysis. The correlation coefficient between *Rp1-D21#4/+* and the *Rp1-D21#4/+; Oy1-N1989/+* double mutants was 0.73 (*p* < 10^-200^) while the correlation coefficient between *Oy1-N1989/+* and *Rp1-D21#4/+; Oy1-N1989/+* was only 0.39 (*p* < 10^-69^). Thus, most of the *Rp1-D21#4*-responsive metabolites are still affected in *Rp1-D21#4/+; Oy1-N1989/+* double mutants. In addition, the *Rp1-D21#4*-induced increased accumulation of SA, SA glucoside, 2,5-DHBA glucoside, and 2,3-DHBA glucoside was also observed for *Rp1-D21#4/+; Oy1-N1989/+* double mutants (Figures 13A-D). Taken together, these findings, along with the insufficiency of SA supplementation to restore *Rp1-D21#4-*induced metabolism in *Rp1-D21#4/+; ics1-1* double mutants suggest that *ics1-1* blocks *Rp1-D21#4-*induced metabolism by a mechanism independent of SA biosynthesis or photosynthetic limitation.

**Figure 13.**
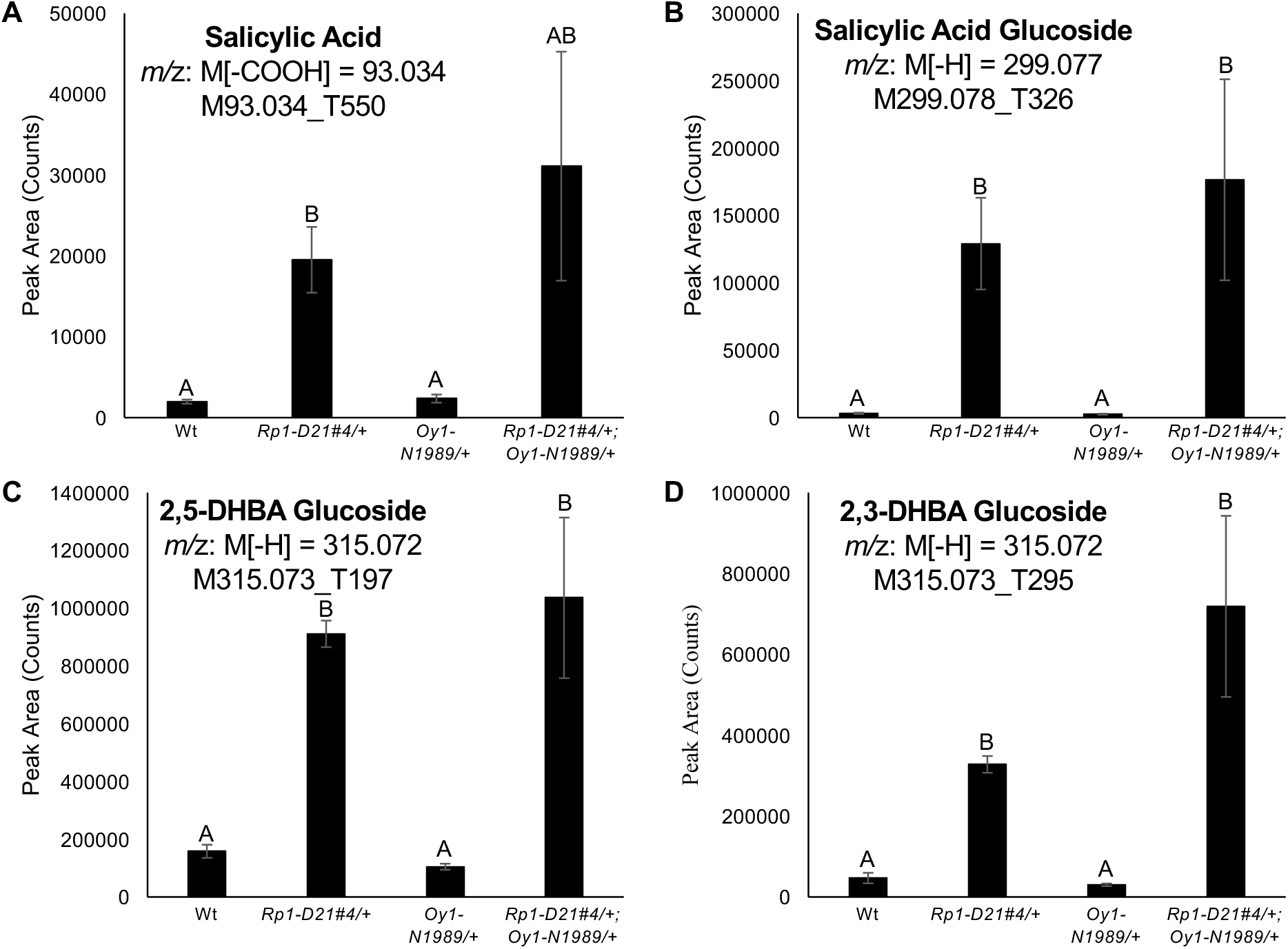
Disruption of photosynthesis does not suppress *Rp1-D21#4*-induced accumulation of SA or SA-derived compounds. Integrated peak areas of (A) salicylic acid, (B) salicylic acid glucoside, (C) 2,5-DHBA glucoside, and (D) 2,3-DHBA glucoside from leaves of Wt, *Rp1-D21#4/+, Oy1-N1989/+*, and *Rp1-D21#4/+; Oy1-N1989/+.* Metabolites were analyzed in negative ionization mode. For salicylic acid, the integrated values are of the major decarboxylation fragment M[-COOH]. For salicylic acid glucoside, 2,5-DHBA glucoside, and 2,3-DHBA glucoside the values are integrations of the parent ion. Data presented are the mean values ± S.E. (n = 4). Letters above bars represent statistically different measurements (paired sample Student’s T-test *p* < 0.05).

## Discussion

In this work we describe a loss-of-function mutant of maize *ics1*. The *ics1-1* allele resulted in a ∼90% reduction in phylloquinone biosynthesis (Figure 6A). While not quite as severe, the *ics1-1* mutants phenotypically more closely resemble *ics1 ics2* double mutants of Arabidopsis than single mutants of either gene (Wildermuth et al., 2001; Gross et al., 2006; Garcion et al., 2008). The two Arabidopsis *ICS* genes do not equally contribute to ICS activity (Garcion et al., 2008), and single mutants of *ics1* are often used for studying the role of ICS in SA production (Wildermuth et al., 2001; Huang et al., 2010). Despite producing only 33% of wild-type phylloquinone levels and about half of wild-type SA levels under control conditions, Arabidopsis *ics1* mutants are morphologically similar to wild-type plants (Garcion et al., 2008) and do not present any stunted growth or leaf yellowing. In contrast, maize *ics1-1*, which produced ∼10% of wild-type phylloquinone levels had severe growth and chlorophyll defects (Figures 5 and 7E). This suggests that plants can sustain normal growth and coloration with reduced phylloquinone and that growth and pigmentation defects manifest when phylloquinone drops below a critical level. The viability of the strong hypofunctional *ics1-1* allele allowed us to examine the role of ICS in phenomena that occur as plants develop and mature, such as the formation of lesions during the hypersensitive response.

### Reduction in photosynthesis was insufficient to block Rp1-D21#4-induced metabolism

Of the photosynthetic parameters that have been measured for both *Oy1-N1989/+* and *ics1-1* mutants, photosynthesis is reduced more by *Oy1-N1989/+* than by *ics1-1* mutants. Compared to wild-type controls, chlorophyll levels were reduced 89% and the maximum efficiency of PSII photochemistry (Fv/Fm) was decreased 41% in *Oy1-N1989/+*:B73 mutants (Khangura et al., 2019; Khangura et al., 2020). These decreases are greater than the 50% drop in chlorophyll (Figure 7e) and 16% drop in Fv/Fm (Figure 7d) in chlorophyll levels and Fv/Fm measured in *ics1-1* mutants. Yet, *Oy1*-mediated loss of photosynthesis did not suppress *Rp1-D21#4*-induced metabolite accumulation (Figure 13 and Supplemental Data Set S5). Thus, energy limitation per se is an unlikely mechanism for suppressing HR-induced metabolism in *Rp1-D21#4;ics1-*1 double mutants. It is possible that the way photosynthesis is altered in *ics1-1* but not *Oy1-N1989* can suppress specialized metabolism. PSII and PSI stoichiometry, the levels of phylloquinone and plastoquinone, and the PSI electron transport rate are unknown in *Oy1-N1989/+* mutants. If these are unaffected by *Oy1-N1989*, it is possible that the shift in PSII and PSI stoichiometry (Figure 8), the loss of phylloquinone or plastoquinone (Figure 6), or the decrease rate of forward electron flow through PSI in *ics1-1* mutants (Figure 7) leads to suppression of and HR-induced metabolism. Inhibiting phylloquinone biosynthesis in *Rp1-D21#4/+* mutants by targeting enzymes in the phylloquinone biosynthetic pathway downstream of ICS (Widhalm and Rhodes 2016) would provide a test of whether a loss of phylloquinone and photosynthetic deficiencies lead to suppression of HR-induced metabolism. In *ics1-1*

### The two known pathways of SA biosynthesis in plants are not independent

In plants, SA can be synthesized from both the ICS-dependent and PAL-dependent pathways (Figure 1). In this work, we have provided evidence for the existence of both pathways in maize. *ics1* was required for SA accumulation (Figure 9), and SA-derivatives can be labeled by ^13^C-phenylalanine (Figures 10 and Supplemental Figure S2). One unexpected finding is that derivatives of the phenylpropanoid pathway and PAL activity both require *ics1* (Table 1 and Figure 11C). This result entangles the two known plant SA biosynthetic pathways and complicates the interpretation of the loss of SA in *ics* mutants in maize and possibly all plants. Because SA can be made from phenylalanine via PAL, and PAL activity was suppressed in the *ics1-1* mutant (Figure 11C), the loss of SA in the *ics1-1* mutant could result, at least in part, from the suppression of PAL. Thus, we cannot regard the requirement of *ics1* for SA accumulation during *Rp1-D21*-mediated HR as evidence that SA is predominantly derived from isochorismate. A retrospective assessment of the literature suggests this is unlikely restricted to maize. For example, previous work in Arabidopsis found that SA was reduced by 90% in *ics1* mutants and 50% in a *PAL1/2/3/4* quadruple mutant (Huang et al., 2010). Similar results were obtained in soybean where SA accumulation was decreased by 90% when ICS was knocked down by virus-induced gene silencing (VIGS) and 95% when PAL was knocked down by VIGS (Shine et al., 2016). If both pathways additively contribute to SA accumulation, then the single pathway knockdowns should add up to 100%, perhaps even less if the loss of one pathway is compensated for by increased flux through the other. Instead, these prior studies and ours indicate that these two pathways require each other in some previously unrecognized manner. Since disruption of *ICS* suppresses PAL activity, our results, and any of the prior genetic experiments are not sufficient to estimate the relative contribution of the two pathways to SA accumulation. Metabolic flux analyses with labeled phenylalanine and labeled isochorismate are required to determine how much each pathway contributes to SA biosynthesis. In our labeling experiment we did not detect phenylalanine-derived labeled free SA. The relative counts of SA were much lower than SA glucoside, 2,5-DHBA glucoside, and 2,3-DHBA glucoside in *Rp1-D21#4/+* mutants (Figure 9). The detection of the more abundant derivatives, and failure to detect labelled SA may be due to hyperconversion of SA to these derivatives. Metabolic flux analyses of SA biosynthesis and catabolism will likely require optimizing SA detection or monitoring of SA-derived products as presented here.

We did not detect the intermediates in the recently described ICS-dependent SA biosynthetic pathway from Arabidopsis (Rekhter et al., 2019; Torrens-Spence et al., 2019) in maize. This failure to detect included *Rp1-D21* mutants which accumulated 9-fold more SA and as much as 99-fold more SA-glucoside. We used similar extraction and chromatography methods as used in the studies of Arabidopsis, so this failure is unlikely to owe to technical variation in the methods. The maize genome encodes a number of genes in the *Gretchen Hagen3* enzyme family, one or more of which may catalyze the same or similar reaction as PBS3, but we did not find any metabolite features with the *m/z* values for isochorismate conjugated with any amino acid that was ICS-dependent. It may be that the PBS3 pathway is taxonomically restricted. PBS3 is a member of a group of the *Gretchen Hagen3* enzyme family which is present in Arabidopsis but not maize and no clear PBS ortholog is encoded by the maize genome (Feng et al., 2014). Consistent with this, *EPS1*, the second step in the Arabidopsis ICS-dependent SA pathway, is restricted to the *Brassicaceae* . This enzyme is unlikely to be strictly required for this pathway as isochorismoyl-glutamate can spontaneously convert to SA (Rekhter et al., 2019; Torrens-Spence et al., 2019). The presence of EPS1 in the *Brassicaceae* may indicate that the ICS-dependent pathway is the predominant route for induced SA synthesis in that plant family. This is further evidenced in the correlation of *ICS* expression and SA accumulation in Arabidopsis (Wildermuth et al., 2001). Further work is needed to demonstrate the generality of this pathway. In tobacco, despite accumulating high levels of SA, *ICS* expression was unchanged following pathogen infection and instead PAL transcripts were increased (Ogawa et al., 2006). In pathogen-infected soybean and *Rp1-D21* mutants, which both accumulate increased levels of SA, ICS transcript accumulation is significantly decreased, and PAL transcripts were increased (Olukolu et al., 2014; Shine et al., 2016) inconsistent with the ICS pathway contributing to the induced accumulation of SA. Thus, in multiple experiments outside the Brassicaceae, including *Rp1-D21*-induced gene expression in maize, *PAL* expression is corelated with SA accumulation and *ICS* expression was not.

### Increased accumulation of SA is not required for Rp1-D21#4-induced HR lesion formation

Accumulation of SA is a hallmark of HR and many autoimmune lesion-forming mutants (Morris et al., 1998; Bruggeman et al., 2015; Radojičić et al., 2018). In this work, we demonstrated that increased SA accumulation is not required for the initiation or progression of lesions caused by *Rp1-D21#4*. The *Rp1-D21#4/+*; *ics1-1* double mutants, which accumulated approximately four-fold less SA than *Rp1-D21#4/+* single mutants (Figure 9), has a similar lesion phenotype as *Rp1-D21#4/+* (Figure 5). In addition, foliar application of SA that increased SA concentration by almost 300-fold compared to mock-treated plants and 17-fold compared to *Rp1-D21/+* (Supplemental Figure S3) did not induce HR-like lesions formation 72 h after treatment. Likewise, SA treatment neither induced the accumulation of HR-induced metabolites, nor rescued the *Rp1-D21*-induced metabolites from the epistatic suppression by *ics1-1* (Table 3 and Supplemental Data Set S3). These findings demonstrate that SA is neither necessary nor sufficient for lesion formation during HR in maize. These findings indicate a difference in signaling between maize and Arabidopsis. In Arabidopsis, conversion of SA to catechol by overexpression of the *NahG* SA hydroxylase suppressed lesion formation and cell death in autoimmune lesion-forming mutants (Weymann et al., 1995; Rate et al., 1999; Brodersen et al., 2005; Huang et al., 2018). HR and cell death can also be triggered in Arabidopsis via supplemental treatment with SA (Dietrich et al., 1994).

SA can be converted to 2,5-DHBA and 2,3-DHBA, which are rapidly glycosylated (Huang et al., 2018). In Arabidopsis, overexpression of *UGT76D1*, which glycosylates 2,5-DHBA and 2,3-DHBA, resulted in increased cell death (Huang et al., 2018), indicating these DHBAs may function in lesion formation during HR. We demonstrated that both 2,5-DHBA glucoside and 2,3-DHBA glucoside accumulated in *Rp1-D21#4/+* mutants (Figure 9) and both compounds accumulated in a concentration-dependent manner after foliar spray with SA (Supplemental Figure S5 and Figure 12). Neither 2,5-DHBA glucoside nor 2,3-DHBA glucoside accumulated in *Rp1-D21#4/+; ics1-1* mutants (Figure 9). Given that none of these treatments altered lesion formation, 2,5-DHBA and 2,3-DHBA were also not required for HR lesion formation in maize. Similarly, we expect that other isochorismate-derived products, like phylloquinone and the pathway intermediate 1,4-dihydroxy-2-naphthoic acid (DHNA) which were proposed to be involved in triggering programmed cell death in Arabidopsis (Brodersen et al., 2005) are also not involved in HR lesion formation in maize.

### Suppression of Rp1-D21#4-induced metabolism in Rp1-D21#4/+; ics1-1 is not dependent on SA

Our untargeted metabolite analysis demonstrated suppression of *Rp1-D21#4*-induced metabolism in *Rp1-D21#4/+; ics1-1* double mutants (Table 2 and Supplemental Table S2). PAL activity was decreased in *ics1-1* mutants (Figure 11C), and of the 46 phenylalanine-derived *Rp1-D21#4*-induced compounds, 37 were reduced in accumulation in the double mutants (Table 1). Just as for SA, these findings indicate that accumulation of these phenylalanine-derived compounds was not required for *Rp1-D21#4*-induced lesion formation. Phenylalanine-derived features accounted for only 5% of those induced by *Rp1-D21#4* and suppressed by the interaction of *Rp1-D21#4* and *ics1-1.* Thus, disrupting *ics1* suppressed multiple *Rp1-D21#4*-induced metabolic pathways. These effects were not rescued by the addition of SA, demonstrating that SA accumulation is neither the trigger nor a modulator of HR-induced metabolic perturbations in this system

Despite evidence that phenylpropanoids are unnecessary for HR-induced lesions, the phenylpropanoid pathway has been linked to HR suppression. Two enzymes likely involved in phenylpropanoid metabolism, a hydroxycinnamoyl transferase paralog and caffeoyl CoA *O*-methyltransferase paralog, physically interact with the *Rp1-D21* protein and suppress HR and lesion formation in response to *Rp1-D21*(Wang et al., 2015; Wang and Balint-Kurti, 2016). Mutations of conserved residues required for the catalytic functions of these enzymes did not interfere with HR suppression indicating that suppression is likely mediated by protein-protein interaction and does not require a phenylalanine-derived product.

That the loss of ICS suppressed most *Rp1-D21#4*-induced metabolites and did not block lesion formation suggests that many of the metabolic changes are consequences of a programmed response rather than the effects of oxidation during cell death. The presence of lesions in the double mutants also demonstrates that HR initiation and progression are not dependent on the soluble metabolites suppressed by *ics1-1*. The 29% of the *Rp1-D21#4*-induced features that are not suppressed by the interaction of *Rp1-D21#4/+*; *ics1-1* contain the remaining candidates for HR induction signaling along with features that accumulate because of cell death. Further experimentation is necessary to determine if these metabolites act in a signaling capacity.

## Methods

### Sequence similarity searches

The maize B73 reference genome (version 4; (Jiao et al., 2017) was downloaded from www.maizegdb.org. The Arabidopsis *ICS1* and *ICS2,* and maize *ics1*, exons were obtained from the Phytozome v13 database (Goodstein et al., 2012). Similar sequences in the maize genome were identified using the amino acid sequences corresponding to each exon as queries and TBLASTN (Gertz et al., 2006) to search for a conceptual translation of ICS in the maize nucleotide genome. Genomic regions matching the exons were determined based on high sequence match and low expectation (E) scores. The conceptual translations corresponding to the full-length protein sequences of Arabidopsis *ICS1*, Arabidopsis *ICS2*, and maize *ics1* were also used for similarity searches of the B73 reference genome (version 4) using TBLASTN and identified the same genomic regions as the single exon searches.

### ICS Phylogenetic analysis

All protein sequences with greater than 50% sequence similarity to Arabidopsis ICS1 (At1G74710) or ICS2 (At1G18870) were downloaded from the Phytozome v13 database (Goodstein et al., 2012). Sequences were aligned using Clustal Omega (Sievers et al., 2011). Sequences deemed partial or incomplete were identified and removed before proceeding with the phylogenic analysis by visualizing the alignments using the constraint-based alignment tool (COBALT) for multiple protein sequences (Papadopoulos and Agarwala, 2007). Phylogenetic trees were generated using IQ-TREE (Trifinopoulos et al., 2016) with the option to output the bootstrap results as a .ufboot file. The IQ-TREE parameters were: 1000 bootstrap alignments; 1000 maximum iterations; a minimum correlation coefficient of 0.99; 1000 replicates of the SH-aLRT branch test; perturbations strength of 0.5; and 100 as the IQ-TREE stopping rule. Phylogenetic Trees were visualized using FigTree v1.4.4 (https://github.com/rambaut/figtree/releases).

### Plant Material and Genotyping

The *ics1-1* mutant allele (mu1039765) was obtained from the UniformMu population (McCarty et al., 2013) in the W22 inbred background as stock UFMu-03865 from the Maize Genetics Co-Op (http://maizecoop.cropsci.uiuc.edu). Genotyping of the *ics1-1* allele was done using three primers, two complementary to *ics1*, ICS1FP 5’-TCTCAGGTACCATTTTGACGA-3’ and ICS1RP 5’-CCCATTCAGGTTCAATCAATGT-3 that span the *Mu* insertion site and one complementary to the *Mu* terminal inverted repeat (TIR) 5**’-**GCCTCCATTTCGTCGAATCCC-3’. All PCR contained 4 µL buffer, 1.6 µL 25 mM MgCl_2_, 0.4 µL 10mM dNTPs, 0.4 µL 10 µM of each primer, 0.1 µL GoTaq® Flexi DNA polymerase (Promega Corporation Ref: M8291), 12.1 µL water, and 1 µL template DNA. The PCR program consisted of an initial 2 min at 95°C followed by 10 cycles of 95°C for 30 s, 65°C for 30 s, decreasing the temperature by 1°C each cycle, and 72°C for 45 s, and then 25 cycles at 95°C for 30 s, 55°C for 30 s, and 72°C for 45 s, and finally 5 minutes at 72°C. Wild-type alleles were detected as a ∼450 bp amplicon using the two *ics1* primers and *ics1-1* alleles were detected as a ∼250 bp amplicon using the *Mu* TIR and ICS1FP primers. After initial genotyping experiments, *ics1-1/ics1-1* homozygous mutants were identified by a yellow leaf phenotype visible by 14d after sowing. Two alleles of *Rp1-D21* were used, the reference *Rp1-D21* allele (Hu et al., 1996) and the *Rp1-D21#4* allele (Karre et al., 2021). Both alleles were maintained as heterozygotes in nearly isogenic B73 backgrounds. *Rp1-D21/+* and *Rp1-D21#4*/+ mutants were identified by 14d after sowing due to lesion formation on the leaves. The *Oy1-N1989* mutants used were maintained as heterozygotes in a nearly isogenic B73 background as previously described (Khangura et al., 2019). *Oy1-N1989/+* mutants were identified by a yellow leaf phenotype after emergence. Material segregating 3:3:1:1 for wild type, *Rp1-D21#4/+*, *ics1-1*, and *Rp1-D21#4/+; ics1-1* was generated by crossing *Rp1-D21#4/+*:*B73* with *ics1-1/+*:*W22* and then crossing an F1 plant without lesions and heterozygous for *ics1-1* (*+/+; ics1-1/+*) with a sibling heterozygous for both *Rp1-D21#4* and *ics1-1* (*Rp1-D21#4/+; ics1-1/+*). Plants segregating 3:1 for wild type:*ics1-1* were generated by self-pollinating an *ics1-1/+* heterozygous plant from the F2 progeny of the above-mentioned sibling mating. Material segregating 1:1:1:1 for wild type, *Rp1-D21#4/+*; *Oy1-N1989/+*, and *Rp1-D21#4/+;Oy1-N1989/+* was generated by crossing an F2 *Rp1-D21#4*/+ mutant from the above-mentioned sibling mating with an *Oy1-N1989/+:B73* mutant.

### Growth Conditions

Material for the phylloquinone analysis, plastoquinone analysis, labeled phenylalanine feeding, all untargeted metabolite profiling, and salicylic acid treatments were grown under greenhouse conditions (mogul-based high-pressure sodium lamps (1000 Watts) on a 16-hour light/8-hour dark cycle with the temperature set at 28°C-day and 20°C-night). Plants were sown in Berger BM2 Seed Germination and Propagation Mix in 36-cell 606 deep plastic flats with one seed per well. Flats were sub-irrigated with fertilizer water as needed, usually every third day after emergence, until tissue collection. All field material was tractor-planted and grown at the Purdue Agronomy Center for Research and Education in West Lafayette, Indiana (40.4700° N, 86.9917° W). Material was planted in 3.84 m rows with interrow spacing of 0.79 m and alley spacing of 0.79 m. Supplemental irrigation was applied as needed, and conventional fertilizer, pest control, and weed control practices for growing maize in Indiana were followed.

### Foliar application and analysis of SA, 2,3-DHBA, and 2,5-DHBA effects

SA, 2,3-dihydroxy benzoic acid (2,3-DHBA), and 2,5-dihydroxy benzoic acid (2,5-DHBA) were dissolved in distilled water and 0.1% V/V Tween-20. 10-day-old plants were spray-treated every 24 hours for three days. For metabolite analyses, the second leaf was collected 24 hours after the last treatment from three plants per replicate.

Calculation of the correlation between SA levels and lesion severity in *Rp1-D21/+* mutants of different genetic backgrounds was performed by reanalysis of the data presented in (Ge et al., 2021). Total and free SA values were obtained by measuring the height of each bar from the graph in Figure 5A (Ge et al., 2021) using a ruler. Correlation analyses of SA accumulation means, and the lesion severity scores in Figure 5B for *Rp1-D21/+* in each genetic background, were performed using the “=correl()” equation in Microsoft Excel.

### Phylloquinone and Plastoquinone Analysis

All steps were conducted in dimmed light to limit the photodegradation of quinones. Approximately 300 mg of flash-frozen ground maize leaf tissue, harvested 17-days after planting, was extracted overnight at 4°C in 3 mL methanol containing 2.7 nmol menaquinone-4 as an internal standard. Phylloquinone and plastoquinone were analyzed directly on an Agilent Infinity 1260 high-performance liquid chromatography (HPLC) system (Agilent Technologies, Palo Alto, CA, USA) coupled with diode array (DAD) and fluorescence (FLD) detectors using a method adapted from previous work (Block et al., 2013, 2014; McCoy et al., 2018). Before analysis, the methanolic extract was filtered using a 0.2 μm polytetrafluoroethylene (PTFE) syringe filter, and 100 μL was directly analyzed on a Zorbax SB-C18 column (4.6 x 250 mm x 5 µm) at 25°C and eluted in isocratic mode at 0.5 mL min^−1^ for 30 minutes with 30% solvent A (60:40 (V/V) isopropanol:hexanes containing 10 mM ammonium acetate) and 70% solvent B (80:20 (V/V) methanol:isopropanol containing 10 mM ammonium acetate). Phylloquinone (retention time 11.8 min) and plastoquinone (23.0 min) were detected fluorometrically (at 238 nm excitation and 426 nm emission and 290 nm excitation and 330 nm emission, respectively). Fluorescence-based detection of phylloquinone was achieved by reducing it in line with a post-column chemical reactor (1.5 x 70 mm) packed with −100 mesh zinc (Sigma-Aldrich) placed between the DAD and FLD modules (van Oostende et al., 2008). Phylloquinone and plastoquinone were quantified relative to external calibration standards and corrected for recovery using a menaquinone-4 internal standard.

### Photosynthetic parameter analysis

The maximum quantum yield of photosystem II (PSII) was measured using a Hansatech pulse amplitude modulated (PAM) fluorescence monitoring system (FMS1) (Norfolk, England). Leaves were dark adapted for 30 min before measurement. Minimum fluorescence (F_o_) was obtained by illuminating with measuring light (≦0.01 µmol m^-2^ s^-1^). Maximum fluorescence (F_m_) for the dark-adapted state was determined with a 0.8-second-long saturating pulse of white light (13000 µmol m^-2^ s^-1^). The actinic light was then switched on, and a second saturating pulse was applied after 6 minutes (F’_m_). Fv/Fm was calculated as 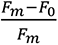, NPQ was calculated as 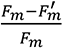, and ΦPSII as 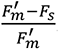. Fraction of reduced PQ-pool (1-qL) was calculated as 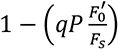, where *qP* = 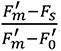. Electron transport rate through PSII (ETR) was calculated as PPFD x 0.84 x 0.5 x ΦPSII. The redox state of P700 was determined by difference absorption spectroscopy of the reaction center chlorophyll P700 using a JTS-10 spectrophotometer (Bio-Logic). The leaves were dark adapted for 30 min before measurement. PSI redox activity was determined by monitoring optical changes at 705 nm during 5 s of illumination with an actinic light peaking at 630 nm (940 µmol m^-2^ s^-1^) and 5 s of dark re-reduction. The amounts of PSI and PSII were determined by difference absorption spectroscopy as described in (McKenzie et al., 2020).

### Untargeted metabolite analysis

Tissue was flash frozen in 15- or 50-mL Falcon tubes and ground using a metal spatula. Soluble metabolites were extracted in 1:1 methanol:water (V/V) at a concentration of 100 mg tissue per 1 mL of extraction solvent. Samples were incubated at 65° C for 2 h and vortex agitated every 30 min. Samples were centrifuged at 10,000 x *g* for 10 min, and 500 µL of supernatant was transferred to a new tube. Samples were dried in a speed vacuum at room temperature and then redissolved in 50 µL 50% aqueous methanol. Samples were sonicated for 15 min and centrifuged at 14,000 x *g* for 10 min. 40 µL of supernatant was transferred to a new tube and stored at -20°C prior to analysis.

Samples were analyzed essentially as described in (Simpson et al., 2021a) using an Agilent 1100 HPLC system (Agilent Technologies, Palo Alto, CA, USA) and a Shimadzu Shim-pack XR-ODS (3.0 × 75 mm × 2.2 µm) separation column and a mobile phase of 0.1% aqueous formic acid (V/V) and solvent B was 0.1% formic acid (V/V) in acetonitrile at a flow rate of 0.6 mL min^-1^. For all analyses, 5 µL of sample was injected, and an initial solvent A:B ratio of 98:2 was held for 1 min. Metabolites were eluted by linear gradients to 94:6 at 5 min, 54:46 at 15 min, 5:95 at 21.5 min, and a 5:95 hold for 2 min. The column was then re-equilibrated by returning to 98:2 over 1 min and holding for 4 min prior to loading the next sample. The column was maintained at 40°C during all steps. Mass spectrometry across a mass-to-charge (*m*/z) range of 80 – 1,100 amu was carried out on the column effluent via negative electrospray ionization (ESI) using an Agilent 6210 time-of-flight mass spectrometer at a capillary voltage of 3.2 kV; N_2_ gas temperature of 325°C; drying gas flow rate of 9 L min^-1^; nebulizer gas pressure of 55 psi; fragmentary voltage of 130 V; skimmer voltage of 45 V; octopole RF of 750 V. Mass accuracy was enhanced by infusing Agilent Reference Mass Correction Solution (G1969-85001) throughout each run. The resulting Agilent MassHunter raw data (.d directories) were visualized and converted to .mzdata files using Agilent MassHunter Qualitative Analysis software (v10.0). Peak picking and retention time alignments were performed using XCMS (Smith et al., 2006) and a custom R script (Supplementary File S1). Peak areas and retention times for all detected features were exported to Microsoft Excel for processing. Linear model analysis was performed using the “lm” function in R (https://www.rdocumentation.org/packages/stats/versions/3.6.2/topics/lm).

All standards were obtained from Sigma-Aldrich. The amino acids used were part of the L-Amino Acids Kit (LAA-21) and salicylic acid was obtained as CAS 69-72-7. Samples were dissolved in 50% methanol and analyzed by LC-MS using the same method as described above. The identity of SA glucoside was determined by the expected spontaneous fragmentation pattern of *m/z*: 299.077 è 137.024 è 93.035 amu and by the response of this feature to foliar application of SA. 2,5-DHBA glucoside and 2,3-DHBA glucoside were identified by the expected spontaneous fragmentation pattern of *m/z*: 315.072 è 151.019 è 109.029 amu and by the response of these features to the foliar application of 2,5-DHBA (CAS 490-79-9) and 2,3-DHBA (CAS 303-38-8) respectively.

### ^13^C-ring labeled phenylalanine feeding

Phenylalanine feedings were performed similarly to the method of (Simpson et al., 2021b). Progeny from a sibling cross of wildtype x *Rp1-D21/+* segregating 1:1 for wildtype: *Rp1-D21/+* were sown in Turface Athletics MVP® (Profile Products LLC, Mfg. Number: BFMVP5004) for approximately 14 d under greenhouse conditions as described above. Three replicates of wildtype and *Rp1-D21/+* were fed 1 mM of phenylalanine (Sigma-Aldrich) or 1 mM ^13^C-ring labeled phenylalanine (Cambridge Isotope Laboratories, Tewksbury, MA. Cat No. CLM-1055) in Murashige and Skoog medium (PlantMedia SKU: 30630058-3). Potting media was washed off roots with water, and whole plants were placed into 120 mL specimen cups containing 30mL feeding media, which was enough to submerge the roots. After 24 h, the basal 2 cm of each plant was harvested and flash frozen in liquid nitrogen. Soluble metabolites were extracted and analyzed using the untargeted LC-MS method as described above.

Labeled features derived from ^13^C-ring labeled phenylalanine were identified as mass feature pairs that coeluted within 5 s and had an *m/z* that increased by 6.02 ± 0.002 amu, as expected from ring-labeled phenylalanine. This was done by duplicating the XCMS output file and running a custom R script (Supplemental File S2) that uses the “fuzzyjoin” package (https://cran.r-project.org/web/packages/fuzzyjoin/index.html) to perform pairwise comparisons of retention times and *m/z* of all detected features. The labeled compound list was trimmed to remove any features annotated as labeled compounds that were also detectable in non-label treatments. This library was then used to annotate phenylalanine-derived metabolites in *Rp1-D21#4/+; ics1-1.* Because metabolite analyses were done at different times, mass features were paired based on a shared *m/z* (± 0.05 amu) and a retention time difference of ± 15 s.

### PAL Assay

Leaf tissue from 15-d-old plants was flash frozen and ground to a fine powder using a mortar and pestle. Immediately after grinding, approximately 400 mg of tissue was transferred to a 1.7 mL Eppendorf tube, and 1.2 mL of extraction buffer (100 mM Tris-HCl pH 8, 10% glycerol, 5 mM dithiothreitol) was added. Tubes were kept on ice until all samples were processed and then centrifuged at 20,000 x *g* for 15 min at 4°C. The supernatants were transferred to new tubes and samples were desalted using Cytiva PD-10 Desalting Columns following the manufacturer’s instructions. Protein concentrations were determined using the Bradford assay with bovine serum albumen as a standard (Bradford, 1976). Each PAL assay contained 190 µL of protein extract and 10 µL of 0.1 M phenylalanine for a total volume of 200 µL. Reactions were incubated for 90 minutes at 37°C. The reactions were quenched with 24 µL of glacial acetic acid and flash frozen in liquid nitrogen. Samples were centrifuged at 14,000 x *g* for 1 h at 4°C. Assay products were quantified by HPLC at 276 nm using cinnamic acid (Eastman Organic Chemicals) as a standard as previously described (Klempien et al., 2012; Wang et al., 2018). PAL activities are reported as the picokatals (pkat) per mg of protein where one pkat is the picomoles of cinnamate produced per second.

## Accession Numbers

The *ics1-1* mutant is available as stock UFMu-03865 from the Maize Genetics Co-Op (http://maizecoop.cropsci.uiuc.edu). Information for mentioned maize genes can be found under the following B73 version4 IDs: *ics1* (Zm000014020220), *Rp1* (Zm00001d023325). Mentioned Arabidopsis gene information can be found under the following AGI accession numbers: *ICS1* (AT1G74710), *ICS2* (AT1G18870), *PAL1* (AT2G37040), *PAL2* (AT3G53260), *PAL3* (AT5G04230), *PAL4* (AT3G10340)

## Supplemental Data

**Supplemental Figure S1.** Retroactive correlation analysis of *Rp1-D21/+* lesion severity and SA accumulation in different genetic backgrounds from data in Ge et al., 2021.

**Supplemental Figure S2.** Incorporation of ^13^C-ing labeled phenylalanine into 2,3- and 2,5-dihydroxybenzoic acid.

**Supplemental Figure S3.** Labeled parent ion of SA was not detected in fed *Rp1-D21/+* mutants.

**Supplemental Figure S4.** Labeled decarboxylation fragment of SA was not detected in fed *Rp1-D21/+* mutants.

**Supplemental Figure S5.** SA and SA-derived compound accumulation in B73 plants treated with different concentrations of SA.

**Supplemental Figure S6.** Leaf damage in plants treated with 5 mM SA.

**Supplemental Figure S7.** Comparison of lesion severity in *Rp1-D21#4/+* mutants and *Rp1-D21#4/+* ; Oy1-N1989 double mutants

**Supplemental Data Set S1.** Processed XCMS output file for the untargeted metabolite analysis of wildtype, *Rp1-D21#4*/+, *ics1-1*, and *Rp1-D21#4/+; ics1-1*

**Supplemental Data Set S2.** All compounds identified as labeled by ^13^C ring-labeled phenylalanine

**Supplemental Data Set S3.** Processed XCMS output file for the untargeted metabolite analysis of B73 plants spray treated with different concentrations of SA.

**Supplemental Data Set S4.** Processed XCMS output file for the untargeted metabolite analysis of wildtype, *Rp1-D21#4*/+, *ics1-1*, and *Rp1-D21#4/+; ics1-1* treated with SA

**Supplemental Data Set S5.** Processed XCMS output file for the untargeted metabolite analysis of wildtype, *Rp1-D21#4*/+, *Oy1-N1989*, and *Rp1-D21#4/+; Oy1-N1989*

**Supplemental File S1.** R script for running XCMS

**Supplemental File S2.** R script for running fuzzyjoin

## Author Contributions and Acknowledgements

RLB, RMM, IMI, JPS, FM-V, CC, JRW, SP, GSJ and BPD designed the experiments. RLB, RMM, IMI, JPS, FM-V, performed the research. RZ, GSJ, and BPD provided genetic stocks and segregating families. RLB, RMM, IMI, JPS, FM-V, JRW, SP, and BPD analyzed the data. RLB, RMM, IMI, JRW, SP, BPD wrote the paper. All authors read and approved the final manuscript.

Metabolite profiling was performed at the Purdue University Metabolite profiling facility at the Bindley Bioscience Center and the authors are profoundly grateful to the staff at the facility and Bruce Cooper, in particular, for his tireless support for excellent data. High performance compute resources were maintained by the Rosen Center of Advanced Computing at Purdue University. We thank Dr. Rajdeep Khangura for the *Oy1-N1989* mutant. All plant material was grown at the Agronomy Center for Research and Education and the Horticulture Greenhouse and the staff of these two amazing facilities are gratefully acknowledged, in particular Jim Beatty and Nathan Deppe, for their leadership and expertise. The work performed was funded a United States Department of Agriculture National Institute of Food and Agriculture predoctoral grant 2019-07171/1022993 to RLB, National Science Foundation grant 1444503 to BPD and GSJ, United States Department of Agriculture National Institute of Food and Agriculture postdoctoral grant 2018-08121/1019231 to JPS, Department of Energy grant DE-SC0020639 to SP, United States Department of Agriculture National Institute of Food and Agriculture predoctoral grant 2018-67011-28032 to R.M.M., United States Department of Agriculture National Institute of Food and Agriculture grant 2021-67013-33779 to J.R.W., United States Department of Energy Office of Science (BER) Grant DE-SC0020368 to CC and BPD.

---

**Supplemental Table S1:**
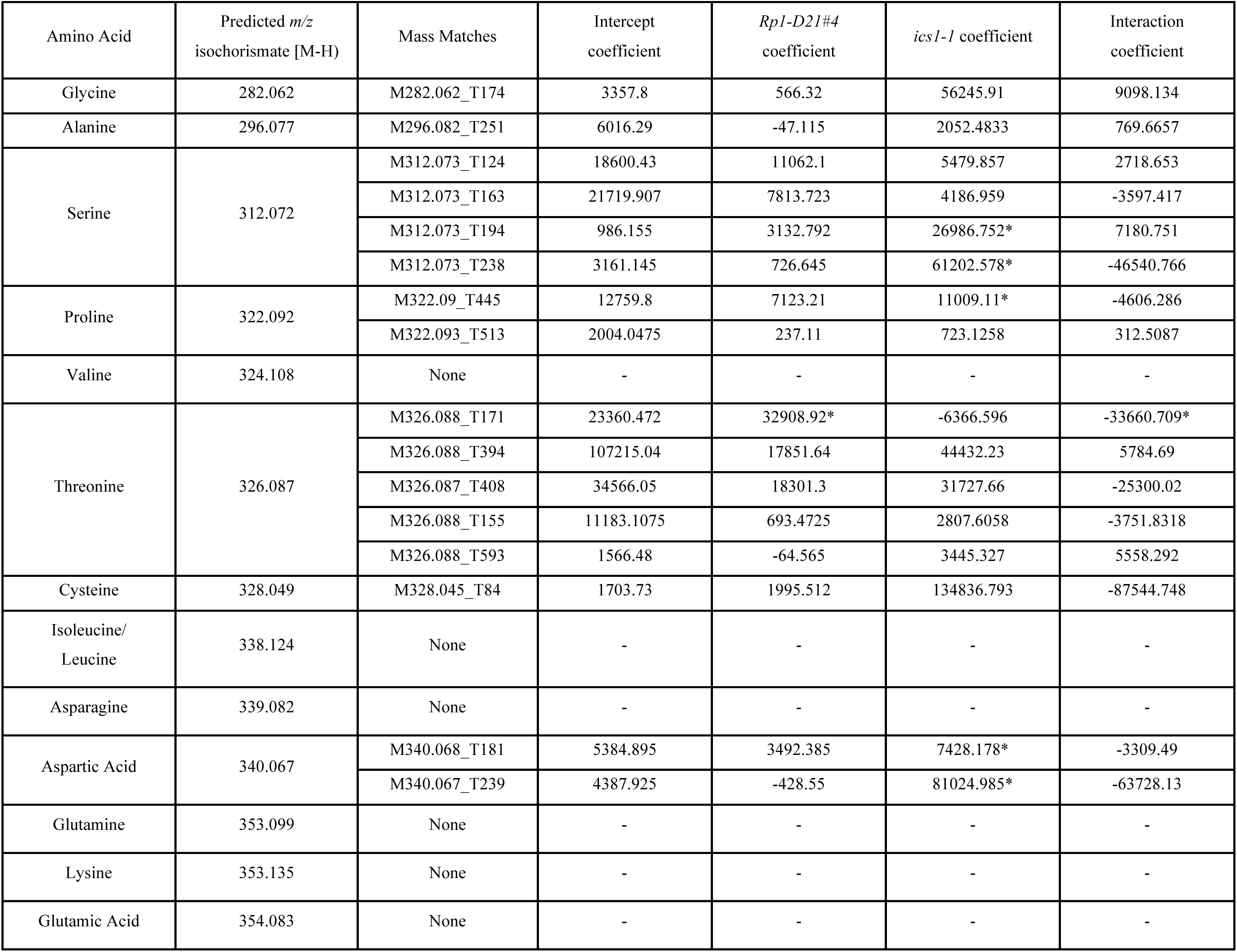
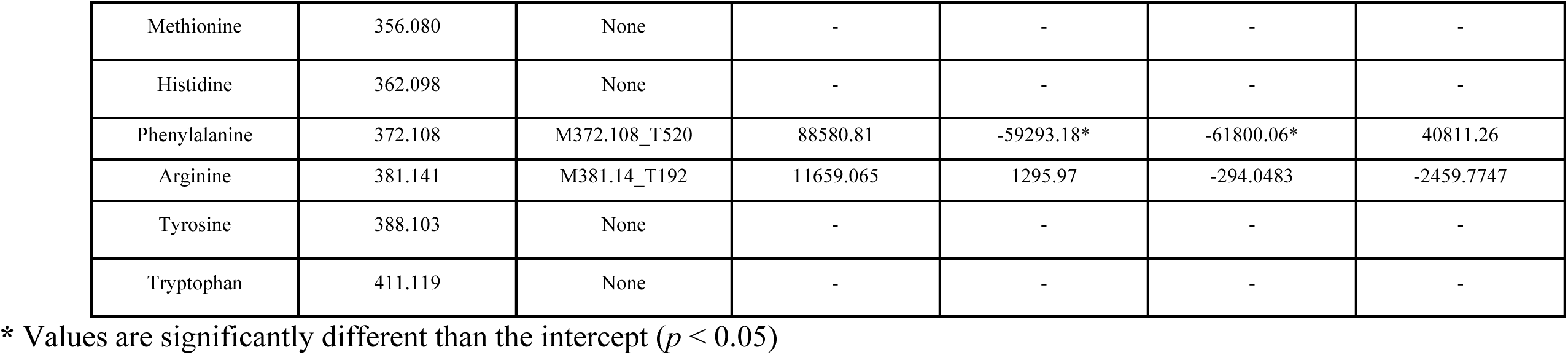
Accumulation of putative isochorismoyl-amino acids in *Rp1-D21#4/+, ics1-1*, and *Rp1-D21#4/+; ics1-1*

**Supplemental Table S2:** Reproducible finding that *Rp1-D21#4*-responsive metabolism is suppressed in *Rp1-D21#4; ics1-1* double mutants

| *Rp1-D21#4*<br>coefficient | *ics1-1*<br>coefficient | Interaction<br>coefficient | *Rp1-D21#4*<br>><br>*ics1-1* | All<sup>a</sup> | FDR<sup>b</sup> | FDR<sup>b</sup><br>*Rp1-D21#4*<sup>c</sup> | FDR<sup>b</sup><br>*ics1-1*<sup>d</sup> | FDR<sup>b</sup><br>*Rp1-D21#4*<br>*ics1-1*<sup>d</sup> | FDR<sup>b</sup><br>*Rp1-D21#4*<br>Interaction<sup>e</sup> | FDR<sup>b</sup><br>*ics1-1*<sup>d</sup><br>Interaction<sup>e</sup> | FDR<sup>b</sup><br>*Rp1-D21#4*<br>*ics1-1*<sup>d</sup><br>Interaction<sup>e</sup> |
| --- | --- | --- | --- | --- | --- | --- | --- | --- | --- | --- | --- |
| + | + | + | Yes | 77 | 10 | 5 | 1 | 1 | 0 | 0 | 0 |
|  |  |  | No | 352 | 211 | 3 | 181 | 3 | 0 | 6 | 0 |
| + | + | - | Yes | 579 | 305 | 303 | 32 | 32 | 227 | 23 | 23 |
|  |  |  | No | 673 | 358 | 51 | 358 | 51 | 26 | 57 | 26 |
| + | - | + | Yes | 219 | 61 | 4 | 57 | 1 | 0 | 0 | 0 |
| + | - | - | Yes | 1375 | 1094 | 873 | 414 | 240 | 648 | 166 | 166 |
| - | + | + | No | 838 | 477 | 99 | 317 | 39 | 25 | 25 | 6 |
| - | + | - | No | 481 | 287 | 4 | 282 | 3 | 0 | 16 | 0 |
| - | - | + | Yes | 1335 | 855 | 433 | 853 | 433 | 250 | 257 | 250 |
|  |  |  | No | 604 | 189 | 184 | 111 | 111 | 134 | 86 | 86 |
| - | - | - | Yes | 99 | 35 | 0 | 30 | 0 | 0 | 0 | 0 |
|  |  |  | No | 42 | 2 | 0 | 0 | 0 | 0 | 0 | 0 |

<sup>a</sup>All 6,676 features, <sup>b</sup>Features with an FDR-adjusted model p value < 0.05, <sup>c</sup>Features whose accumulation was significantly affected by *Rp1-D21#4* (p < 0.05), <sup>d</sup>Features whose accumulation was significantly affected by *ics1-1* (p < 0.05), <sup>e</sup>Features whose accumulation was significantly affected by the interaction of *Rp1-D21#4* and *ics1-1* (p < 0.05). Columns with multiple headings represent features who meet all criteria as described above

